# Systematic Evaluation of Nasal Immune Cell Sampling and Antigen-specific T cell Detection using Cryopreserved Nasal Swabs

**DOI:** 10.64898/2026.08.27.747456

**Authors:** W. Huisman, M.A. Power, N. le Bert, E. Mitsi, N. Verleur, L.A. King, D.W. de Vos, G.E. Loe-Sack-Sioe, H.R. Wagstaffe, R. S. Hagedoorn, I. Satti, C. H. GeurtsvanKessel, A. Marchant, P. Pannus, M.H.M. Heemskerk, H. McShane, R.S. Thwaites, C. Chiu, G.H. Groeneveld, D. Ferreira, R.D. de Vries, S.P. Jochems

## Abstract

The upper respiratory tract is a key entry point for pathogens, yet local tissue-resident memory T cells (Trm) remain underexplored compared to peripheral blood. We systematically compared nasal curettes and 8 different swab types for immune cell collection, assessing yield, operator variability, and T-cell phenotypes across the three turbinates and nasopharynx. The use of flocked swabs yielded higher immune cell numbers while being similarly tolerated, especially with reduced sampling duration. Nasal Trm subsets were consistent across the turbinates, whereas nasopharyngeal Trm displayed a more recently recruited phenotype. Multiple cryopreservation media were evaluated and all demonstrated high viability after thawing. Antigen-specificity was assessed using the activation induced marker (AIM) assay, peptide–HLA tetramers and bulk TCR-sequencing following expansion. Notably, influenza-specific T cell frequencies were reliably detected by AIM and correlated between fresh and cryopreserved nasal samples. Downregulation of the CD3/TCR complex was observed in nasal samples. These findings establish a robust approach for nasal Trm profiling, demonstrating that cryopreservation preserves functional antigen-specific T cells. This work enables centralized, minimally invasive nasal T cell analysis for multicenter studies, including mucosal vaccination trials and controlled human infection models.

## Introduction

The human upper respiratory tract (URT) serves as the primary entry point for respiratory pathogens, emphasizing the critical role of local mucosal immunity in antiviral defense. Multiple cell types are involved in mucosal immunity, among which tissue-resident memory T cells (Trm) are essential mediators capable of quickly detecting and responding to viral (re-)infection, helping to limit pathogen transmission^1–4^. Recent human studies have demonstrated that the nasal mucosa harbors antigen-specific Trm populations that differ from circulating T cells, reinforcing the need to study human mucosal tissues to accurately understand responses to respiratory infections and vaccine-induced mucosal T cell immunity^5–7^. Traditionally, however, investigations of virus-specific T cell responses have relied on peripheral blood samples, which may not fully reflect local mucosal immunity^8^.

In peripheral blood, several established methodologies are available to quantify and characterize antigen-specific T cell responses, including ELISPOT, activation-induced marker (AIM) assays, cytokine release assays, peptide-HLA tetramer staining, TCR sequencing, and *in vitro* expansion approaches^5–10^. Together, these methods have substantially advanced our understanding of antiviral and vaccine-induced T cell immunity in humans. However, their applicability to nasal mucosal samples remains largely unknown.

Minimally invasive sampling techniques such as flocked nasal swabs and nasal curettage are currently used as practical tools to sample nasal-derived immune cells without significant discomfort, allowing for repeated sampling^7, 9^. Different sampling methods and sites within the nasal cavity have been studied, including the inferior turbinate (IT), middle turbinate (MT), and superior turbinate (ST), as well as the nasopharynx (NP), each with distinct protocols^5–8^. However, no systematic evaluation has compared the different anatomical sites and sampling protocols to determine which site provides the highest yield with the lowest variability. In addition, freshly collected nasal material is currently regarded as the most suitable for phenotypic profiling of Trm and performing antigen-specific assays due to the low number of cells that can be collected from nasal samples ^5–7, 9^; however, relying on fresh samples poses logistical challenges for large-scale multicenter studies, highlighting the urgent need for cryopreservation methods that retain the functionality of Trm in nasal samples.

In this study, as part of the MUSICC consortium, we systematically evaluated the various parameters required for studying antigen-specific nasal cells. We assessed optimal sampling protocols, comparing different cryopreservation media, with the goal of the viability and antigen-specific functionality of nasal Trm. We demonstrated that antigen-specific nasal Trm were detectable from the same donors both fresh and from frozen samples using AIM assays.

## Methods

### Study participants

All participants provided written informed consent in accordance with the Declaration of Helsinki. Nasal curette samples were obtained as part of the TINO trial (clinicialtrials.gov:NCT06039527), a prospective cohort study investigating the mechanisms and underlying cause of T cell decline in the upper-respiratory tract of the aging population that was conducted at the Leiden University Medical Center (LUMC, Netherlands). Healthy young adults (18-30 years) and older adults with and without frailty (>65 years) were included. Participants with recent respiratory infection (<2 weeks), recent vaccination (<2 months), or any airway related co-morbidity were excluded from participating. Ethical approval was obtained from the Medical Ethical Committee Leiden-Den Haag-Delft (NL77841.058.21). For sampling technique comparisons (curette versus flocked swabs), samples were obtained from healthy voluntary donor services from the Leiden University Voluntary Donor Service (LuVDS; protocol LuVDS25.014). This protocol describes the use of these samples for optimization and comparing techniques only. Additionally, nasal flocked swabs were obtained as part of the RESPECCT trial, a randomized controlled trial to look at the effect of RSV and Pneumococcal co-infection conducted at Oxford Vaccine Group, (Oxford University, UK), or from a study approved by the SIngHealth Centralized Institutional Review Board (Singapore; CIRB/F 2021/2014). To compare the phenotype of anatomical locations, flocked swabs were collected from the LuVDS and SENTINEL study (Surveillance of rEspiratory viruses iN healThcare and anImal workers in the NethErLands) conducted at Erasmus Medical Center. Ethical approval was obtained from the Medical Ethical Committee Erasmus Medisch Centrum Rotterdam (NL86800.078.24). Peripheral Blood Mononuclear cells (PBMCs) were obtained as part of the SENTINEL study and were isolated by standard Ficoll-Isopaque separation and stored in the vapor phase of liquid nitrogen.

### Nasal sample collection and cell isolation

#### Nasal curettes

Nasal cells from the inferior turbinate that were collected using curettes were isolated as described previously^9^. Briefly, curettes (ASL Rhino-Pro©, Arlington Scientific) were used to scrape a small collection of cells from the inferior turbinate. Two curettes per nostril were used and stored in a 15 mL Falcon tube placed on ice containing 8mL phosphate-buffered saline (PBS), 0.5% heat-inactivated fetal bovine serum (hiFBS, PAN Biotech, United Kingdom), and 2.5 mM ethylenediaminetetraacetic acid (EDTA). Cells were dislodged from the curettes by repeated pipetting with collection medium from the same tube. Cells were spun down (450 x g for 5 minutes) and pellets from the two 15 mL Falcon tubes were pooled, resuspended in 1 mL Cryostor CS10 (Stemcell), and aliquoted into two cryovials. Step-wise freezing was performed by using pre-chilled Mr. Frosties before long-term cryopreservation in a liquid nitrogen tank.

#### Nasal swabs

Nasal cells from different anatomical locations of the upper respiratory tract were isolated similarly as previously described^5, 7^ . To compare yield of nasal-derived cells from different types of swabs, the inferior turbinate (IT) was swabbed by rotating for 10 seconds using 8 different types of swabs (**Supplementary Table 1, Supplementary Figure 1**). Swabs were transferred to a cryovial containing 1 mL Cryostore CS10 and cryopreserved according to the manufacturer’s protocol. To compare different anatomical locations, flocked swabs (COPAN; ref:220252) were inserted into the IT, mid-turbinate (MT), superior turbinate (ST), or nasopharynx (NP) of the donors and rotated for 10 seconds or 30 seconds. A single swab from each nostril was then processed immediately (fresh) or transferred to a cryovial containing 1 mL Cryostor CS10, Bambanker, CryoABC, or 10% DMSO in hiFBS and cryopreserved according to the manufacturer’s protocol. Swabs that were immediately processed were collected in 2 mL isolation medium; Roswell Park Memorial Institute (RPMI) 1640 (Gibco, United Kingdom) containing 2% hiFBS, 100 µg/mL streptomycin (Gibco, United Kingdom) and 100 U/mL penicillin (Gibco, United Kingdom) (Pen/Strep) and 1.5mM Dithiothreitol (DTT; Merck). DTT was added fresh just before use. Swabs were then incubated for 30 minutes at 37°C. Flocked swabs were then taken out using sterile forceps and rinsed twice with fresh isolation medium. The single cell suspension was then centrifuged for 7 minutes at 400g.

### Scoring the tolerability of nasal sampling

Following nasal curettage and nasal flocked swabbing, some of the participants rated on a 5-point modified Likert scale how much pain, discomfort, and lacrimation were experienced for each procedure, as described previously^9^.

### Nasal phenotyping using spectral flow cytometry

Cryopreserved nasal curettes or flocked swabs were thawed in a 37°C water bath and transferred to a 15 mL Falcon tube, and washed by slowly adding 4 mL pre-warmed thawing medium (RPMI 1640 Medium (Gibco, United Kingdom), 20% hiFCS, 100 µg/mL streptomycin, and 100 U/mL penicillin. Nasal cells from frozen flocked swabs were then additionally processed with DTT as described above. Freshly processed or thawed nasal samples were then taken up in 180 μL PBS and transferred to a V-bottom 96-well Nunclon™ Delta Surface plate (Thermo Fisher Scientific, Denmark), and cells were centrifuged for 7 minutes at 400g. After centrifuging and removing the supernatant, cells were resuspended and stained in 50 µl for 15 minutes at RT with LIVE/DEAD™ Blue fixable Blue Dead Cell Stain (Invitrogen, USA) solution at 1:500 dilution mixed with anti-human Fc Receptor (FcR) Binding Inhibitor (Invitrogen, USA). Then extracellular staining was performed by adding 50 µl of 2x concentrated mix containing either a small phenotyping panel (n=9; #1-#9) or large phenotyping panel (n=33) of monoclonal antibodies directed against extracellular proteins (Supplementary **Table 2**) prepared in FACS (PBS containing 2 % (v/v) BSA, 2 mM EDTA)) buffer with 10 % (v/v) BD Horizon™ Brilliant Stain Buffer Plus (BD Biosciences, USA) for 15 minutes at room temperature (RT). For the analysis of tissue-resident marker retention after cryopreservation using different freezing media, the same procedure described above was used, with a small phenotyping panel (n=10) of monoclonal antibodies directed against extracellular proteins (**Supplementary Table 3**). For analysis of the median fluorescence intensity of TCRαβ, the same staining procedure was used using the small phenotyping panel (**Supplementary Table 2**), replacing aCD11c-PE with aTCRαβ-PE at 1:100 (BD). After extracellular staining, cells were washed twice with FACS buffer and then resuspended in 200 µL of FACS buffer for acquisition. Cells were acquired with fluidics boost on a 5-laser Aurora Cytometer (Cytek Biosciences, Inc., USA). Reference controls were made using either AbC™ Total Antibody Compensation Beads (Invitrogen, USA) or PBMCs. An additional unstained control, gated on monocytes (FSC/SSC), was used for unmixing.

### Analysis of reactive T-cells with activation-induced marker (AIM) assays using flow cytometry

To study the effect of cryopreservation on the reactivity of T cells, paired cryopreserved and fresh nasal cells from flocked swabs were processed as described above and stimulated with Staphylococcal Enterotoxin B (SEB, Sigma-Aldrich) or peptides derived from Influenza. After thawing, processed nasal samples were divided into two wells of a round-bottom 96-well plate each containing 100 µL of T-cell medium composed of RPMI 1640, L-Glutamine, and 25 mM HEPES (Capricorn Scientific). The medium was supplemented with 10% heat-inactivated human pooled serum (hiHPS, Sanquin), 100 IU/mL penicillin, and 100 IU/mL streptomycin. Nasal-derived cells were either stimulated with 100 µl of T-cell medium containing 200ng/mL SEB, a peptide pool (1 µg/mL) containing immunodominant 15-mer epitopes (n=169) derived from internal influenza proteins (Flu-nonHA)^11^, or as a negative control, cells were stimulated with an equimolar amount of dimethyl sulfoxide (DMSO). Similarly, cryopreserved

PBMCs were used to study the reactivity of Influenza-specific T-cells. For PBMCs, thawing was performed using IMDM (Lonza, Belgium) supplemented with 10% hiFBS, 100 IU/ml penicillin, 100 μg/ml streptomycin, and 2 mM L-glutamine. Thawed PBMC were treated with 50 U/ml Benzonase (Merck) for 30 min at 37 °C before use in AIM. For nasal phenotyping, nasal-derived samples stimulated with SEB, Flu-nonHA, or medium control were stained for surface markers at 4 °C for 30 minutes (**Supplementary Table 4**). PBMC samples stimulated with Flu-nonHA peptide pools or DMSO control were resuspended and stained in 50 µl for 15 min at RT with LIVE/DEAD™ Fixable Aqua Dead Cell staining solution or LIVE/DEAD™ Fixable Far Red Dead Cell staining at 1:100 dilution, respectively. Cells were then washed with 125 μL of FACS buffer twice and centrifuged at 400g for 5 min. Then, extracellular staining was performed for nasal samples by adding 50 µl of surface marker mix (**Supplementary Table 5**). After extracellular staining, cells were washed twice with FACS buffer and then resuspended in 150 µL of FACS buffer for acquisition. A lower limit of detection (LLOD) of 0.001% was used to enable reproducible detection of AIM+ cells within the CD4+ or CD8+ T cell gates.

### Detection of antigen-specific T cells using peptideMHC-tetramers

Paired PBMCs and nasal cells taken from the IT using nasal curettes were, prior to peptideMHC-tetramer staining, incubated for 15 min with 50 μM of reversible protein kinase inhibitor (PKI) desatinib (Sigma-Aldrich) at 37°C to lower the T-cell receptor (TCR)/pMHC interaction affinity threshold required for antigen-specific cell staining^12^. Cells were then incubated with a [2x] concentrated pool of peptideMHC-tetramers (**Supplementary Table 6**) for 15 min at 37°C. To differentiate Spike-specific responses from Membrane/Nucleus and other (e.g. ORF1, ORF3, and polyproteins) (MNO), we designed combinatorial labelling of Spike-derived pMHC-tetramers in PE and APC, and MNO-derived pMHC-tetramers in PE and BUV615. Cells were then washed with FACS buffer, centrifuged at 400g for 7 min and then, as described above, stained for LIVE/DEAD™ Blue fixable Blue Dead Cell Stain and extracellular markers **(Table 2**: #1-#7), excluding CD14-APC and CD11c-PE.

### T-cell receptor identification

TCRαβ sequences of T cell populations were identified as previously described^8, 10^. In short, total RNA (10 μL) was extracted using the ReliaPrep RNA cell Miniprep system (Promega) from cryopreserved (Cryostor CS10) nasal samples collected using nasal curettes longitudinally 3 months apart (Timepoint A and B) or from cultured and expanded nasal-derived T cells. Nasal-derived T cells were non-specifically expanded using 0.1 x 10^6^ irradiated (35 Gy) allogeneic feeders as described previously for PBMCs^13^. Preamplification of the Template Switching Oligo (TSO) cDNA was performed on samples containing RNA from 500 or fewer cells. Barcoded TCR PCR product was generated in two rounds of PCR. In the second PCR, the first purified PCR product was used to include a two-sided six-nucleotide barcode sequence that allows for discrimination between TCRs from different T cell populations. PCR products of different T cell populations were pooled, after which TCR sequences were identified by NovaSeq (GenomeScan). NovaSeq data were analyzed using MiXCR software (v3.0.13) to determine the Vα and Vβ families and CDR3 regions using annotation to the IMGT library (http://www.imgt.org; v6). CDR3 sequences with ≤0.1% of total reads, that were non-functional or occurred in all samples were excluded from the analysis. Overlap of TCR sequences was calculated by dividing the number of shared CDR3α or CDR3β sequences between timepoint A and B or *in vitro* expanded by the sum of all CDR3α or CDR3β sequences from timepoint A.

## Results

### Short-duration nasal flocked swabbing generates higher yields and is better tolerated than curettage

To detect antigen-specific T cells from samples obtained from the upper respiratory tract (URT), maximizing cell yields should be a high priority, while maintaining tolerability for participants. Tolerability and yield of sample collection from the inferior turbinate (**Figure 1A**) were compared between performing two curettes from one nostril (RhinoPro©) and one flocked swab (COPAN©-503CS01) from the other nostril (n=6). The flocked swabs were collected by a single trained operator by rotations at the inferior turbinate (IT) for 30 seconds for each participant, as described previously^7^. The same operator collected the curettes from the other side by scraping the inferior turbinate 3-4 times^9^.

**Figure 1.**
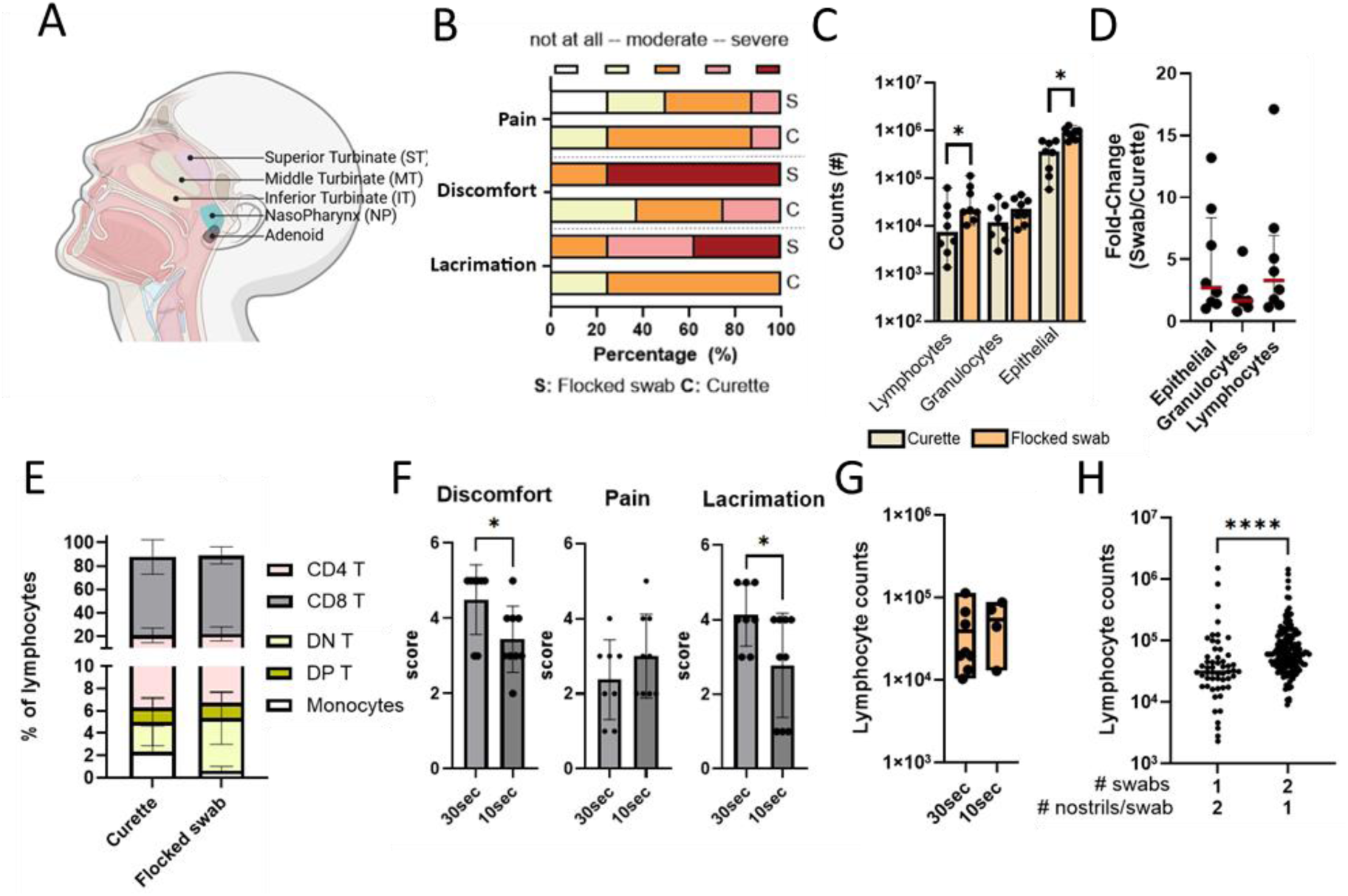
Yield and tolerability of nasal curettes and nasal flocked swabs. Nasal cells can be collected using different techniques; here, the yield and tolerability of two nasal curettes (RhinoPro©) were compared to one nasal flocked swab (COPAN®). **A**) Cross-section of the upper-respiratory tract, showing the different anatomical locations of sampling. **B**) General tolerability was assessed in 8 individuals, where nasal curettes (C) and flocked swabs (S) were collected from the inferior turbinate, randomized across left and right nostrils. Flocked swabs were inserted for 30 seconds, and 3 to 4 scrapes were performed per nasal curette. Lacrimation, discomfort, and pain were assessed. **C**) Yield is shown directly after sampling (fresh) of 8 different donors using flocked swabs or nasal curettes. Cells were acquired using a spectral flow cytometer. **D**) Fold-changes in yield between flocked swabs and nasal curettes are depicted for each cell type. **E**) Cellular composition of live lymphocytes from freshly collected flocked swabs and nasal curettes of 8 donors. **F)** General tolerability was compared between flocked swabs inserted for 10seconds or 30 seconds. **G**) Lymphocyte counts are depicted for flocked swabs that were inserted and rotated at the inferior turbinate for 30 seconds (n=8 donors) versus 10 seconds (n=4 donors). **H**) Yield of live lymphocytes was compared of nasal cells that were collected using one nasal flocked swab (COPAN®) for both sides of the nose (1x 2 nostrils) or collected using two nasal flocked swabs, one for each side. Statistical differences were assessed with Wilcoxon matched-pairs signed rank tests with Bonferonni correction for multiple testing (C and E), Mann-Whitney U tests (F and G) or unpaired T test (H). *P < 0.05; **P < 0.01; ***P < 0.001; ****P < 0.0001

Although the flocked swab was not considered painful, severe lacrimation and discomfort were reported due to the long sampling duration (30 seconds), which was not the case with the curettes (**Figure 1B**). However, flocked swabs yielded significantly more lymphocytes, granulocytes, and epithelial cells than curettes, with a similar immune cell composition (**Figure 1C, 1D, and 1E**). To test whether tolerability for flocked swabs could be increased, sampling duration was reduced to 10 seconds. This significantly reduced discomfort and lacrimation (**Figure 1F and Supplementary Figure 2A**). Sampling for 10 seconds yielded a median of 57,903 lymphocytes, which did not differ significantly from the number of lymphocytes obtained with 30-second rotations (median 20,673, **Figure 1G**). In addition, the T cell composition was comparable between the sampling durations (**Supplementary Figure 2B**). To test how sampling both nostrils could lead to enhanced cell recovery, both sides were sampled using one swab (n=167) per side (2×1; median 60,000 after combining both swabs) compared with using a single swab (n=54) for both sides (1×2; median 31,100), indicating saturation of the swab when sampling both sides with one swab (**Figure 1H**).

We then compared 8 different swab types for sample yield, including flocked and non-flocked swabs with or without a flexible tip (**Figure 2A and Supplementary Figure 1**). Flocked swabs yielded significantly (*P=0.02*) higher relative abundances of lymphocytes and demonstrated improved cell viability (**Figure 2B-2E**). Among the flocked swabs, the COPAN® 503CS01 (H) and M10141 (F) performed best, exhibiting superior cellular yield, viability, CD4:CD8 ratios, and frequencies of T cells with a tissue-resident memory phenotype (**Figure 2B-2E and Supplementary Figure 3**). Of these, only the 503CS01 was available with a flexible tip, enabling sampling from different anatomical regions. Although the M10141 showed similarly strong performance, it was also associated with the highest reported discomfort and pain levels during sampling (**Supplementary Figure 3C**). Therefore, the COPAN® 503CS01 swab was selected for use throughout the study.

**Figure 2.**
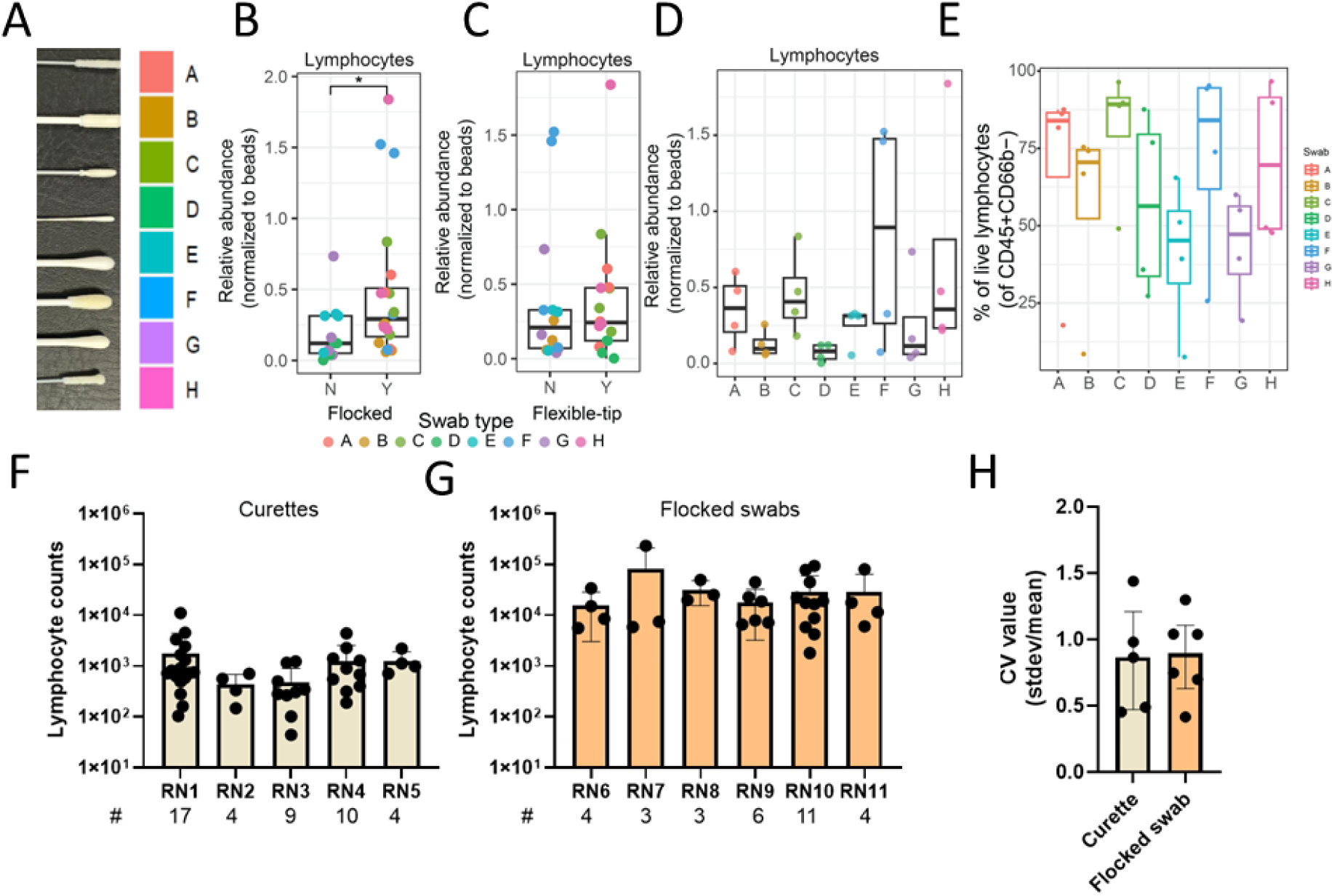
**Cellular yield of different nasal swab types and assessment of operator variability between techniques**. Nasal cells can be collected using different techniques, here operator variability of using two nasal curretes (RhinoPro©) were compared to one nasal flocked swab (COPAN®503CS01). Additionally different swab types were compared for their yield. Swab type H corresponded to the commonly used COPAN®503CS01. **A**) Eight swab types were evaluated for cellular yield and tolerability following 10 seconds of swabbing of the inferior turbinate. Swabs from two manufacturers (Puritan and Copan), including both flocked (Y) and non-flocked (N) designs as well as swabs with flexible (Y) or rigid tips (N), were compared. **B and C**) Yield of live lymphocytes (excluding granulocytes) was compared between (**B**) flocked and non-flocked swabs and (**C**) swabs with flexible versus rigid tips. **D**) Yield of live lymphocytes (excluding granulocytes) is shown for all eight swab types. **E**) Percentage of live lymhpocytes (excluding granulcotes) is shown of total lymphocytes for all eight swab types. **F and G**) Yield of lymphocytes is shown per research nurse (RN) of cryopreserved nasal cells collected using nasal curettes (**F**) or nasal flocked swabs (**G**). Cells were acquired using a spectral flow-cytometer. Number of different donors is depicted below each bar. **H**) Operator variability is shown of operators using curettes and were compared to variability when using flocked swabs. Variability (CV) was calculated by dividing the standard deviation by the mean of lymphocyte counts. Statistical differences were assessed with Mann-Whitney U tests (B, C and H). *P < 0.05; **P < 0.01; ***P < 0.001; ****P < 0.0001

Next, we assessed the variability between operators who collected nasal curettes or flocked swabs to see whether one method was easier to implement across different sites and operators **(Figure 2F and 2G)**. Lymphocyte yield varied considerably between operators for both approaches. Across different operators, the coefficient of variation (CV) of the median recovered lymphocyte counts was similar for flocked swabs and curettes (0.36 versus 0.40). Within operators, the CVs of recovered lymphocytes across different samples were also similar between nasal curettes (median 0.86) and flocked swabs (median 0.89) (**Figure 2H** and **Supplementary Figure 4**).

Overall, we found that nasal flocked swabs yielded more cells than curettes, while showing similar variability between operators. Reducing the flocked swab sampling duration to 10 seconds reduced discomfort without compromising yield, while sampling both nostrils with two separate swabs increased the sampling yield compared to using a single swab for both sides. The commonly used COPAN® 503CS01 was identified as optimal swab for recovery of T cells.

### Comparison between immune-cell phenotypes obtained from swabs at different locations

To investigate which nasal anatomical location would be the best source of Trms, flocked swabs were collected from the IT, MT, ST, and NP (**Figure 1A**). Freshly collected swabs taken by two independent operators in different study centra showed that NP swabs yielded most lymphocytes (median:28,481 cells; *P=0.0067*), whereas MT swabs had lowest recovery (median:3190 cells) (**Figure 3A**). NP swabs recovered T and B cell populations that differed from the swabs collected from the three turbinates. NP swabs contained significantly higher frequencies of B cells, while CD8^+^ T cell frequencies were decreased **(Figure 3B)**. The increased B cell frequencies in the NP swabs may be derived from the adenoids, as demonstrated by a recent study^6^. No differences in immune composition between turbinate locations were observed. In-depth phenotyping was performed on a subset (n=36) of fresh nasal samples to further investigate immunophenotypic differences between anatomical locations. In total, 27 different immune cell populations were manually gated (**Supplementary Figure 5**) and overlaid on a UMAP (**Figure 3C, Supplementary Figure 6**). A distinct immune signature was identified from immune cells derived from NP by UMAP, while cells from all three turbinates had a similar phenotype (**Figure 3D**). NP swabs showed an increase in B cells that were CD11c^-^ and a shift in both CD4^+^ and CD8^+^ T cells from CD103^+^ Trms to CD103^-^ Trms relative to turbinate swabs (**Figure 3E**). Most of the CD4^+^ Trm that were CD69^+^ from the NP were CD127^+^ and/or CD27^+^CD28^+^, while turbinate-derived CD4 T cells were PD1^+^. Similarly, the CD8^+^ Trms from NP lacked CD161 and mostly PD1 (**Supplementary Figure 7B**), explaining the difference in immune signatures as identified by the density UMAPs.

**Figure 3.**
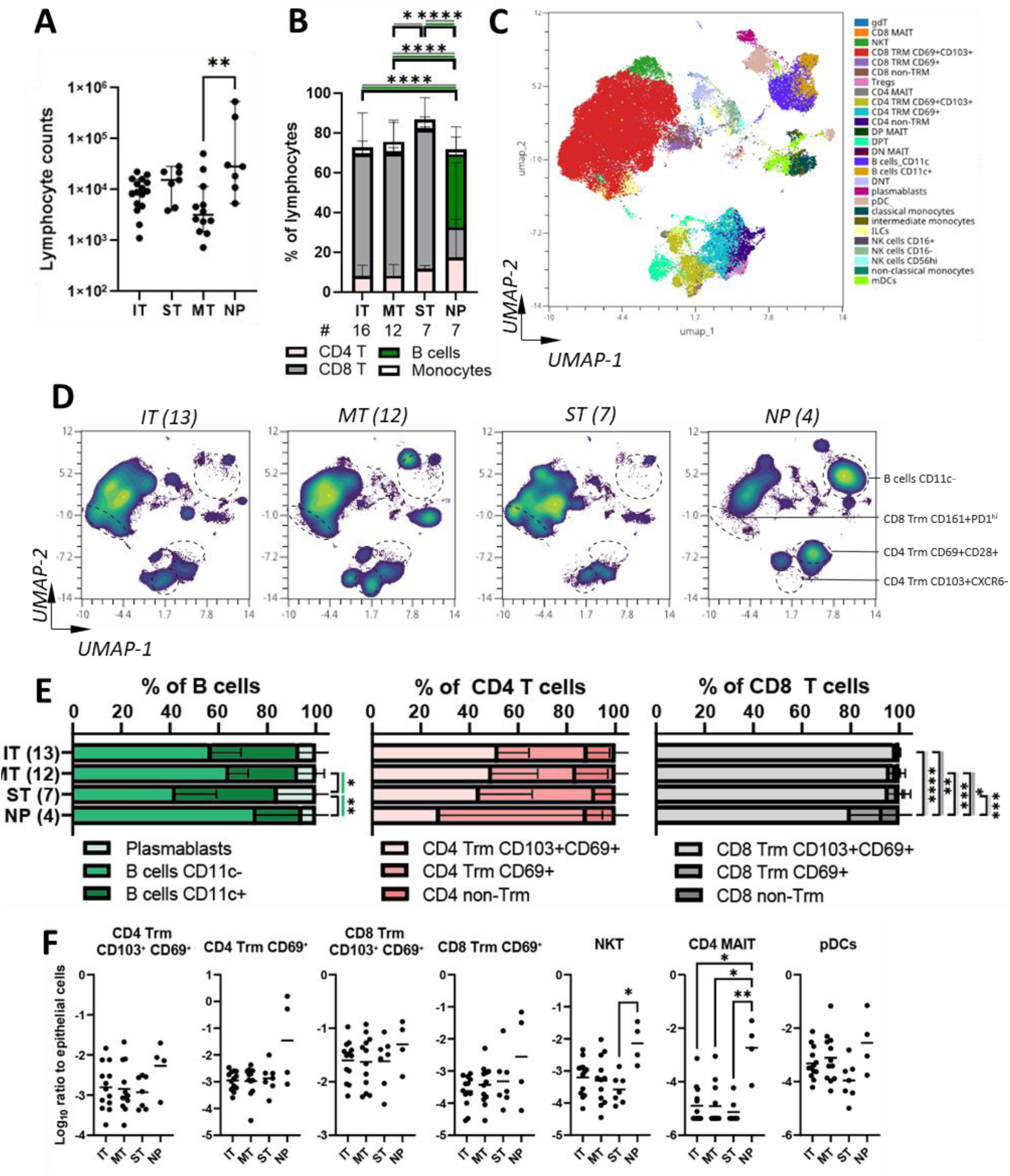
Distinct immune-cell phenotype and composition of nasal swabs from the upper respiratory tract. Nasal swabbing was performed at different anatomical locations (inferior turbinate (IT), middle turbinate (MT), superior turbinate (ST), or nasopharynx (NP)) of the upper respiratory tract. Sampling was performed by two independent operators. **A**) Yield of lymphocytes, (CD45+CDccb-) was compared between freshly collected IT, MT, ST and NP flocked swabs. **B**) Cellular composition of live lymphocytes was compared between freshly collected IT, MT, ST, and NP flocked swabs. **C**) UMAP of 84k nasal-derived lymphocytes, 21k per group, to cluster cells based on 30 different markers. Manually gated populations are overlaid. **D**) Density plots for UMAPs for each sampling location, 21k lymphocytes per group. **E**) Cellular composition of different subsets of B cells, CD4 T cells, and CD8 T cells was compared between freshly collected IT, MT, ST, and NP flocked swabs. **F**) Normalized abundances (log10 ratio relative to epithelial cells) of live nasal immune cell populations. Individual values and box plots are shown. Samples in which a given cell population was undetectable were assigned the lowest detectable value observed for that population. Statistical differences were assessed with 2-way ANOVA with Tukey’s correction for multiple testing (B, and E) or Kruskal-Wallis tests with Dunn’s correction for multiple testing were performed (A and F). *P < 0.05; **P < 0.01; ***P < 0.001; ****P < 0.0001

Further analyses revealed that the abundance (ratio to epithelial cells) of the different manually gated immune cell populations was largely different between the NP and the three turbinates, highlighting the similarity in phenotype and abundances between the turbinates (**Figure 3F**). Flocked swabs from the NP contained higher abundance of different innate lymphocyte populations such as NK cells, NKT cells, γδ T cells and CD4^+^ MAIT cells (**Supplementary Figure 7**).

To summarize, the three turbinates yielded cells with similar phenotypes and counts of immune cell populations with a epithelial-associated Trm phenotype. The inferior and superior turbinate provided higher yield than the middle turbinate, likely because it is harder to correctly sample. In contrast, cells from the NP showed significant phenotypic shifts, indicative of more recently recruited phenotypes compared to the turbinates.

### Serum-free cryopreservation media enable whole flocked swab storage for later analysis

To reduce technical variability and improve sample logistics in particular for multi-centre studies, we next assessed the possibility of cryopreserving whole flocked swabs and detect antigen-specific T cells. First, the number of live lymphocytes was directly compared after processing swabs and cryopreserving half the sample in commercial Cryostor 10 (CS10) medium (**Figure 4A; top panel, Figure 4B**). Lymphocyte counts decreased with a median of 51% (from 55.883 cells to 30.429 lymphocytes) upon cryopreservation of isolated cells, with a small drop of 5% viability (from 90% median to 85%) (**Figure 4B and 4C**). Next, we compared cell recovery and viability of nasal flocked swabs that were directly cryopreserved in four different freeze media: Cryostor 10 (CS10), Bambanker (Bam), CryoABC (Cryo), and FBS supplemented with 10% DMSO (FBS) **(Figure 4A: bottom panel**). Nasal flocked swabs obtained from 12 healthy donors, randomized across the left and right side of the nose for different freeze media (n=6 each), were cryopreserved whole using staged freezing in a freezing container (CryoABC, FBS, or CS10) or directly at −80°C (Bam) **(Figure 4A)**. High viability (median: 83%) of T cells was observed across all freezing media **(Supplementary Figure 8B**). Viable lymphocytes and T cells were recovered from swabs stored in CS10 (median: 1.714 cells), CryoABC (median: 1.439 cells), Bambanker, (median: 580 cells) and FBS (median: 233 cells) (**Figure 4D and 4E, Supplementary Figure 8C and 8D**. The CD4/CD8 composition after thawing was widely conserved across all freeze media **(Figure 4F)**. The CD4/CD8 ratio after thawing was conserved across all freezing media **(Figure 4F)**. Additionally, the overall Trm phenotype, characterized by the expression of CD103, CXCR6, CD49a, CD69 and PD-1, was consistent across freezing media **(Supplementary Figure 8E and 8F)**. In summary, all commercially available serum-free cryopreservation media demonstrated high cell viability after processing.

**Figure 4.**
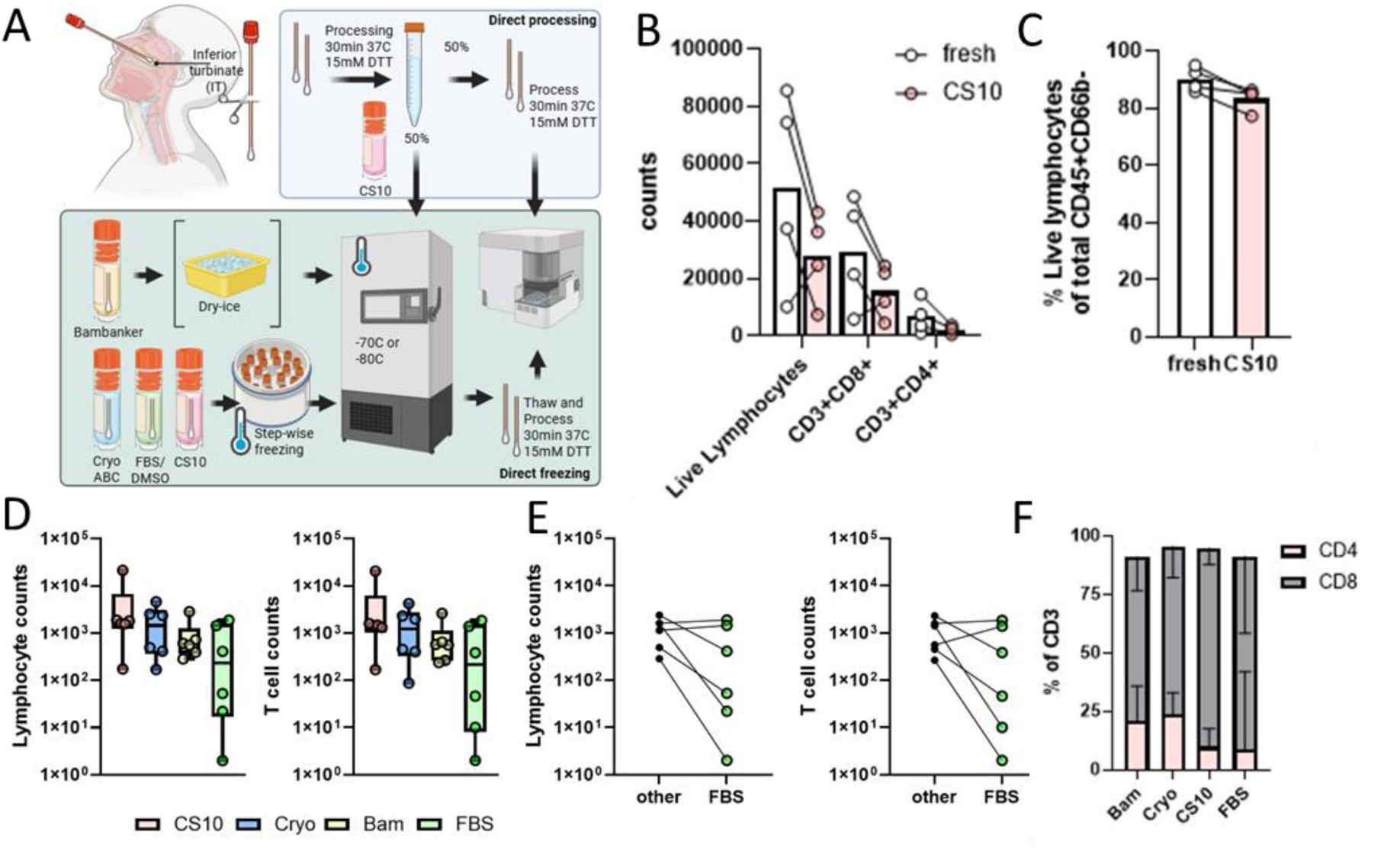
Evaluation of cryopreservation media in nasal flocked swab samples. Nasal sampling using flocked swabs was performed at the inferior turbinate (IT) and immediately stored in one of the following freezing media: Cryostor CS10 (CS10), CryoABC (Cryo), Bambunker (Bam) and 10%FBS/DMSO (FBS)), and then cryopreserved according to the manufacturer’s protocol. For a subset of samples, both swabs from the same donor were first processed, pooled, and 50% was cryopreserved while the other 50% was recorded fresh. In total, samples from 12 healthy donors, randomized across left and right nostrils for different freezing media (n=6 each), were assessed. **A**) Overview of sample processing for different freezing media. The top panel shows the setup for comparing fresh versus cryopreserved cell yields and viability. The bottom panel shows the experimental setup used to compare yields across different freezing media. **B**) Shown are lymphocyte yields and T cell subsets recorded fresh (white) or after cryopreservation with Cryostor CS10 (pink). **C**) The viability of lymphocytes is shown immediately after collection (fresh) or after cryopreservation with Cryostor CS10 (pink). **D)** Total counts of live lymphocytes CD45^+^ CDccb^-^ and live T cells. Individual datapoints and box plots are shown for counts after thawing for every freezing media. **E**) Counts of paired samples are shown, whereby FBS was compared to paired samples cryopreserved with Bambanker, CryoABC, or Cryostor CS10. These are indicated as other. **F**) Percentages of CD4^+^ T cells and CD8^+^ T cells from cryopreserved IT flocked swabs were compared between freezing media. Statistical differences were assessed with the Kruskal-Wallis test with Dunn’s correction for multiple testing (D) or paired t-tests (E). *P < 0.05.

### Detection of antigen-specific cells using cryopreserved nasal flocked swabs

Current protocols for the detection of antigen-specific nasal-derived T cells rely on analysis of fresh material. Peripheral blood studies indicate that cryopreservation may alter T-cell functionality^14^ while largely preserving antigen-specific response magnitude^15^. Whether nasal-derived immune cells remain similarly functional after cryopreservation is unknown. Therefore, cryopreserved and fresh cells from the same individuals (n=14) were obtained, and samples from 4 individuals were stimulated *in vitro* with Staphylococcal Enterotoxin B (SEB), followed by a comparison of reactivity using AIM. Across all 4 donors, cryopreservation followed by SEB stimulation resulted in a gradual reduction of AIM⁺ T cells, with median decreases of 4.6% for CD8⁺ T cells (from 19.8% to 15.2% AIM^+^ after freezing) and 6.6% for CD4⁺ T cells (from 14% to 7.4%) (**Figure 5A and 5B, Supplementary Figure 9A, 9B and Supplementary Figure 10**). While frequencies of responding cells were decreased, the inter-donor variability remained visible upon freezing. Next, T cells from paired fresh and cryopreserved nasal flocked swabs (n=10) were stimulated with a previously described Influenza mega peptide pool (n=169) covering immunogenic 15-mer peptides from all proteins except haemagglutinin (HA) to look at the Influenza-specific T cell response^11^. Influenza-specific CD8⁺ T cells were detected in the same donors both fresh as well as after cryopreservation (n=8 out of 10), as indicated by upregulation of CD137 and/or CD25 (**Figure 5C**). Across all donors with detectable Influenza-specific CD8^+^ T cells, cryopreservation followed by peptide stimulation resulted in a gradual reduction of AIM⁺ T cells, with a median decrease from 2.2% to 1% influenza-specific CD8^+^ T cells after freezing. The frequency of AIM^+^ CD8^+^ T cells before cryopreservation correlated significantly (Rho=0.9, *P=0.0002*) with the frequency of AIM^+^ CD8 T cells after cryopreservation/thawing, indicating maintenance of the magnitude of the response (**Figure 5D**). The number of CD4^+^ T cells per condition (fresh: median 105; frozen: median 95 cells) was too low to support comparisons between fresh and cryopreserved samples. However, when donors were selected with more than 250 live CD4^+^ T cells, either fresh or cryopreserved, we were able to detect influenza-specific CD4^+^ T cells in 3 out of 5 donors (**Supplementary Figure 9C**). In contrast, when looking at the systemic response, influenza CD4^+^ T cells targeting non-HA peptides were more dominant than the influenza CD8^+^ T cells (**Supplementary Figure 9D**).

**Figure 5.**
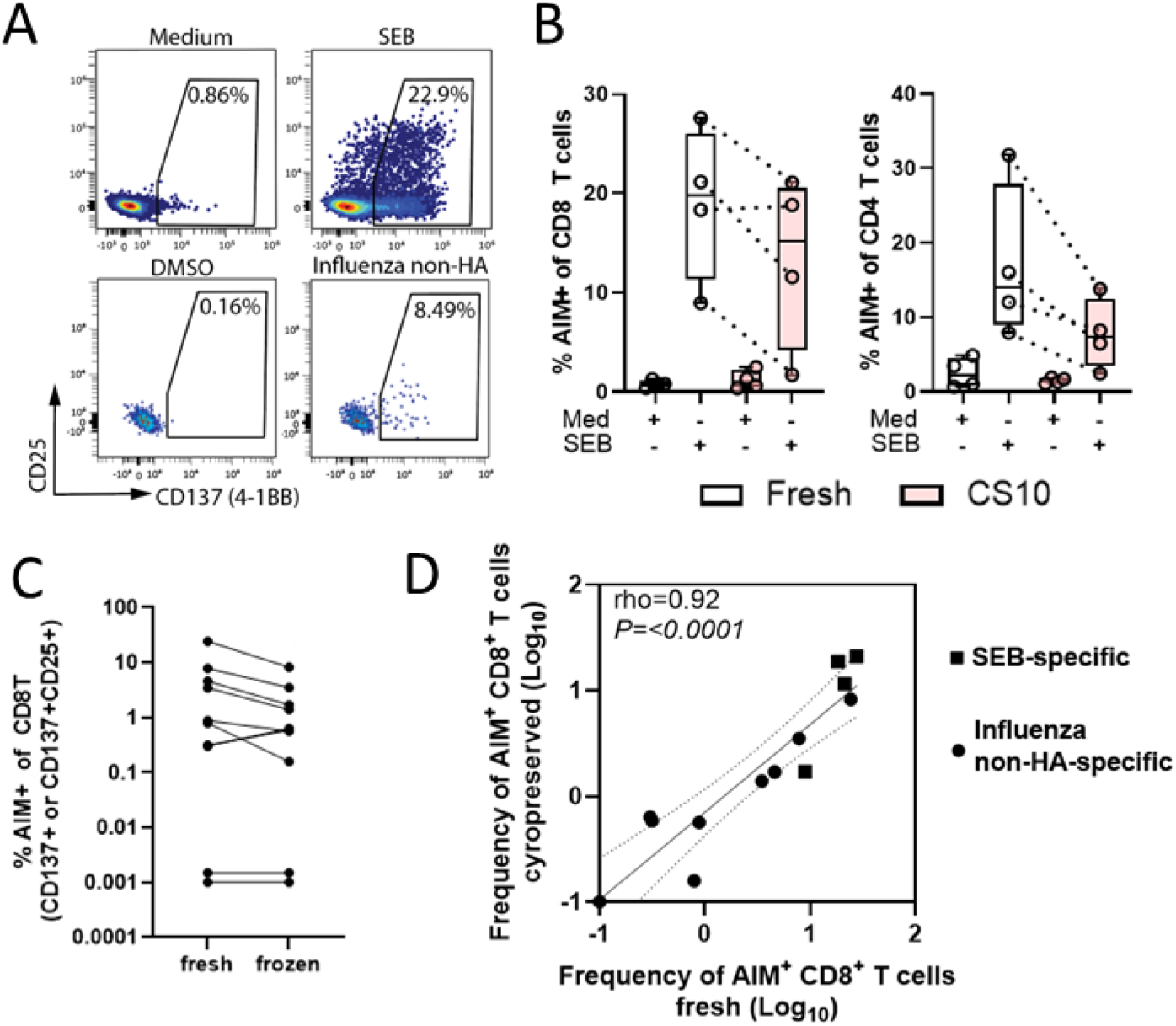
Detection of antigen-specific T cells using cryopreserved nasal flocked swabs. Nasal swabbing using flocked swabs was performed at the inferior turbinate (IT) and either immediately put into freeze media (Cryostor CS10) and frozen according manufacturers protocol or immediately processed and recorded. In total, samples from 14 healthy donors were used to assess differences between fresh and frozen (n=4 for SEB and n=10 for Influenza-specific responses). **A**) Representative flow cytometry plot of CD8^+^ T cells showing upregulation of activation induced markers (AIM) after medium and SEB stimulation or DMSO and an overlapping pool of Influenza peptides (non-HA). **B**) Shown are floating boxplots of activated CD8^+^ or CD4^+^ T cells that were stimulated with SEB immediately after collection (white) or after cryopreservation with cryostor CS10 (pink). Percentage of cells that show AIM^+^ was assessed using expression of either CD137 and/or CD25 for CD8^+^ T cells and CD137 and/or CD25/OX40 for CD4^+^ T cells. **C**) Shown are dotplots of activated CD8^+^ T cells that were stimulated fresh or after cryopreservation using CS10 with an overlapping pool of Influenza peptides (non-HA). Data shown are medium (DMSO) subtracted. Percentage of AIM^+^ CD8^+^ T cells represent the sum of cells that express either CD137 and/or CD25. Samples without AIM^+^ events were plotted at 0.001. **D**) Pearson correlation plot for the frequency of AIM^+^ CD8^+^ T cells (Log_10_ transformed) cryopreserved against the frequency of AIM^+^ CD8^+^ T cells measured fresh (Log_10_ transformed). Responses against the overlapping pool of Influenza peptides (non-HA) are shown as circles and responses against SEB are shown as squares. Samples without AIM+ events were plotted at −1.

Our findings demonstrate that AIM-based functional analysis of cryopreserved nasal flocked swab samples is feasible and enables reliable detection of antigen-specific CD8^+^ and CD4^+^ T cells. Even though absolute frequencies of responding cells were about 50% lower both for SEB and antigen-specific stimulations, the relative magnitude between donors was preserved and freezing did not affect the ability to detect influenza pool-responding donors.

### TCR downregulation and clonal bias following expansion limit quantification of antigen-specific T cells from nasal samples using tetramers or sequencing

We next explored whether direct *ex vivo* quantification of antigen-specific cells using pMHC-tetramers has potential for use in nasal samples. We used paired PBMCs and nasal samples from SARS-CoV-2 infected individuals, 1 month after symptom onset. SARS-CoV-2-specific T cells could be detected in PBMCs using pMHC-tetramers. While pMHC-tetramer-positive cells were detectable, a lower median fluorescence intensity (MFI) of pMHC-tetramer binding was observed in nasal-derived T cells (**Figure 6A**). We hypothesized that intrinsically lower expression of the CD3: T cell Receptor (TCR) complex of nasal-derived T cells might lead to decreased binding of pMHC-tetramers. Therefore, we assessed the median fluorescence intensity (MFI) of the TCR, CD3, and co-receptors CD4/CD8 on nasal-derived T cells from IT, MT, and NP, and compared these to PBMCs. For CD8^+^ T cells, the MFI of CD3, CD8, and TCR was significantly lower in nasal-derived T cells compared to PBMC-derived T cells (**Figure 6B**). Similar decreases in TCR and CD3 levels were observed for nasal CD4^+^ T cells, although the MFI of the CD4 co-receptor was significantly higher on nasal-derived T cells. (**Figure 6B and Supplementary Figure 11A and 11B**). Next, we assessed whether there were differences in MFI of the TCR of different nasal T cell subsets. For CD4^+^ T cells, the TCR MFI was increased in non-TRM versus TRM, while for CD8+ T cells, there was no association with subsets **(Supplementary Figure 11C and 11D**).

**Figure 6.**
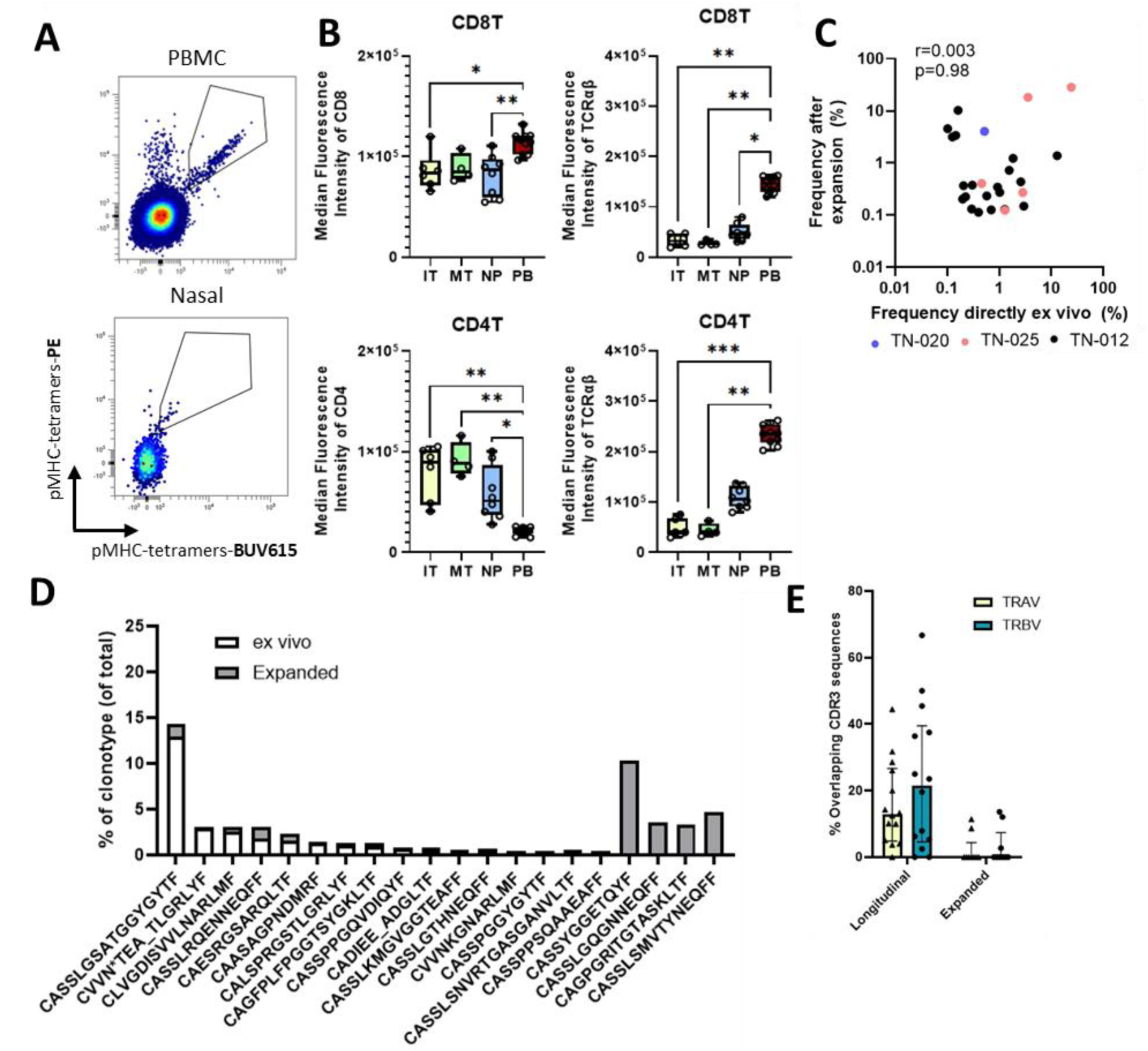
TCR downregulation and clonal bias following expansion limit detection of antigen-specific T cells from nasal samples. Nasal mucosa (nasal curettes) was investigated for presence of antigen-specific T cells of 4 SARS-CoV-2 infected individuals. **A**) CD8^+^ T cells (CD3⁺CD8⁺CD4⁻CD5c⁻) from peripheral blood and nasal curette samples of two individuals one month after SARS-CoV-2 infection, specific for Membrane Nucleus and ORF (MNO) protein– derived peptides, detected using pools of peptide–HLA complexes covering common HLA molecules. **B**) Median Fluorescent Intensity (MFI) of CD8 or CD4 co-receptors and TCRαβ are shown of CD4^+^ T cells and CD8^+^ T cells from PBMCs (red) and different anatomical locations of the upper-respiratory tract, obtained using flocked swabs. **C**) Pearson correlation plot for shared clonotypes. Frequencies of each CDR3β clonotype directly ex vivo (x-axis) was plotted against the frequency of the shared clonotype after in vitro expansion (y-axis). **D**) Representative example of the frequencies of each shared CDR3β clonotype before expansion (white) and after expansion (grey) from one donor (TN-012). **E**) TCR-sequencing was performed for nasal curette samples from individuals where curettage was performed at two consecutive timepoints (A-B, 3 months in between) and some nasal curette samples were non-specifically expanded in vitro. Shown is the percentage of overlap between CDR3α (TRAV) and CDR3β (TRBV)-sequences on timepoint A and B (Longitudinal) and between ex vivo and after expansion (Expanded). Statistical differences were assessed with Kruskal Wallis test with Dunn’s correction for multiple testing (B). *P < 0.05; **P < 0.01; ***P < 0.001

To investigate whether non-specific expansion of nasal material enables relative quantification of the antigen-specific T cell pool while overcoming the limited cell yield, we performed TCR-sequencing on T cells from nasal material before and after 14 days of non-specific expansion *in vitro*. As a comparator, TCR overlap between longitudinally collected nasal material at two time points (3 months) was compared, representing the *in vivo* stability of the nasal T cell pool. We found almost no overlap in TCR clonotypes pre and post expansion. Moreover, clonotypes that were identified after *in vitro* expansion did not correlate with the frequencies measured directly *ex vivo* (**Figure 6C and 6D),** indicating that expansion led to skewing of T cell repertoires and would affect detection. In contrast, a median of 22% (IQR: 4.6-39.5%) of CDR3β and 13% (IQR: 4.8%-26.7%) of CDR3α sequences of TCRs could be found back in longitudinally collected *ex vivo* samples, showing the relative stability of T cell clones in the human nasal mucosa (**Figure 6E**).

These data show that *in vitro* expansion of nasal material leads to a skewed T cell proliferation with only a limited number of clones expanding. Moreover, decreased TCR expression on nasal T cells results in a decrease in pMHC-tetramer binding, making quantification of antigen-specific T cells difficult when using conventional pMHC-tetramers.

## Discussion

Here, we systematically evaluated how sampling location and protocols affect nasal T cell recovery and phenotype. We also assessed the effect of cryopreservation media on nasal-derived T cells and functionality. Finally, we compared different methods for the detection of antigen-specific T cells in samples from the nasal mucosa.

A major challenge in upper respiratory tract mucosal immunology is the lack of standardized approaches for sample collection, anatomical location and processing. Studies differ in both sampling methodology, including swabs, curettage, brushes and lavage-based techniques, and in downstream workflows such as fresh versus cryopreserved analysis, making comparisons across studies difficult^6–9, 16–20^. We demonstrated that both curettage and swabbing could be implemented consistently following basic training, with comparable operator-dependent variability across methods. However, flocked swabs yielded significantly higher numbers of lymphocytes and, in our experience, required less technical expertise and fewer training days than curettage. Anatomically, we showed that the turbinates closely resemble each other and operators should therefore focus on the easiest accessible turbinate to avoid low yields, which is the inferior turbinate.

Related to the lack of standardized approaches is the lack of a study that investigated the use of cryopreservation of nasal-derived samples, thereby relying on freshly processed samples^7^. Although cryopreservation would greatly simplify logistics for large-scale and multicenter studies, its impact on antigen-specific T cell measurements in nasal samples has remained largely unexplored. Here, we demonstrate that antigen-specific T cells can be detected in cryopreserved nasal samples without the need for fresh processing or PBMC co-cultures^5, 9, 21, 22^. While cryopreservation resulted in modest reductions in viability and about 50% reduced functional responses compared with fresh samples, the relative ranking of donor responses was preserved, consistent with observations for cryopreserved PBMCs stimulated with anti-CD3/CD28 antibodies^23^. Importantly, stimulation with influenza A virus peptide pools identified virus-specific CD8^+^ T cells in 8 out of 10 donors both fresh and cryopreserved, demonstrating the biological relevance of the assay. These findings suggest that cryopreservation is a practical and analytically robust approach for nasal immune monitoring including the detection of antigen-specific T cells, substantially reducing logistical barriers for vaccine trials and CHIM studies, particularly in multicenter and resource-limited settings^21^.

We furthermore showed that choosing an anatomical sampling site determines the composition and functionality of the immune cells recovered. Immune profiling revealed nearly identical phenotypes across the three turbinates, whereas NP sampling yielded distinct results, with more B cells, higher ratio of CD4^+^ to CD8^+^ T cells, and overall higher lymphocyte yields. A recent study has shown that B cells from the NP are predominantly germinal-center B cells, with CD4^+^ T cells skewed toward T follicular helper phenotypes^6, 20, 24^. Our findings confirm this, and by performing in-depth phenotyping we further identified differences between Trms obtained from the NP and those from the turbinates. For example, NP-derived cells include Trms co-expressing CD27 and CD28 while lacking CD161 and PD-1, and displayed higher surface TCR:CD3, a phenotype consistent with recently recruited or less terminally differentiated cells rather than the tissue-adapted turbinate Trm profile (CD161+, PD-1+, lower surface TCR expression). Trm from the turbinates showed increased expression of CD103, which interacts with epithelium, and longitudinal TCR-sequencing revealed persistence of clonotypes over time, indicating that the Trm compartment in the human nasal turbinates is more stable.

Several considerations govern how antigen-specific T cells can be detected in nasal samples. The reduced TCR expression on turbinate-derived T cells most likely resulted in low pMHC-tetramer staining intensity, which explains the difficulty of directly detecting antigen-specific cells in the turbinates using conventional pMHC-tetramers^25^. As an alternative, dextramers or spheromers with increased binding affinity might provide higher resolution for detecting these low-TCR-expressing antigen-specific T cells in nasal samples^26, 27^. Another approach is to first expand nasal-derived T cells non-specifically *in vitro*. Although repertoire skewing is a concern, T cells from peripheral blood have been shown to maintain their clonotype hierarchy after expansion^25^. For nasal-derived samples, the pattern was markedly different. Only 3 of 9 donors showed any overlapping clonotypes after expansion. Moreover, even in these donors, clonotype frequencies before and after expansion did not correlate, indicating that expansion substantially reshaped clonal hierarchy rather than faithfully preserving the starting repertoire. Potential reasons for this discrepancy may include: (1) unequal proliferation of individual T cell clones during expansion or (2) a highly polyclonal repertoire, resulting in the majority of unique clonotypes being randomly distributed to pre-expansion and post-expansion cultures. Importantly, non-specific expansion can be used to detect antigen-specific T cells, but it is not reliable for estimating their true frequency or clonal composition.

This study has limitations that should be acknowledged. We also did not collected repeated samples in a short period to assess repeatability of the findings, which we are aiming to do in the future. Although AIM detected antigen-specific T cells after cryopreservation, we did not directly compare assays with or without autologous APC, nor did we benchmark against IFN-γ ELISPOT, which remains the most commonly used T cell assay in the field, or CRA.. Conducting head-to-head assay comparisons in nasal-derived samples will be essential, as AIM has been shown in some studies using peripheral blood to be more sensitive than cytokine-based assays^28, 29^, while ELISpot remains a benchmark for low-frequency responses and has outperformed ICS in some comparisons^30, 31^. AIM, ELISpot, and ICS/CRA each capture overlapping but distinct aspects of antigen-specific T cell responses in PBMC^32^, further underscoring the need to perform such comparisons in mucosal samples.

Our findings establish and validate a practical, scalable workflow for cryopreservation of nasal samples and downstream immune cell analysis, directly filling existing methodological gaps in human mucosal immunology, especially for multi-centre studies. As more intranasal vaccines are developed, using cryopreserved nasal endpoints collected over time, before and after challenge infection or vaccination, will help speed up the discovery of protective mucosal correlates and guide the design of next-generation vaccines.

## Acknowledgements

We thank all research nurses and/or physicians who were responsible for sample collection, and additionally, all healthy volunteers for taking part in this study. The authors gratefully acknowledge dr. Cilia Pothast from Leiden University Medical Center for the generation of peptide-MHC-tetramers and furthermore the flow cytometry core facility (FCF) at LUMC, Leiden, the Netherlands, for technical support regarding spectral flow cytometry. We would also like to thank Dr. Ricardo da Silva and Prof. dr. Alessandro Sette from the La Jolla Institute for Immunology for their influenza mega-peptide pools. This project has received co-funding from the Coalition for Epidemic Preparedness Innovations (CEPI) and the European Union’s Horizon Europe Program as part of the MUSiCC consortium (*Grant reference: PRJ-7359*). Views and opinions expressed are those of the author(s) only and do not necessarily reflect those of CEPI and the European Union or Horizon Europe. Neither the European Union nor the granting authority can be held responsible for them. This work was further supported by an NWO grant (OCENW.KLEIN.461) and a ZonMw grant (10430072110011) to SPJ.

## Author contributions

Conceptualization and study design: WH, MAP, NB, EM, HRW, IS, CHG, AM, PP, MHMH, HM, RST, GHG, CC, DF, RDV, SPJ; Sample collection and processing: WH, MAP, DWV, LAK, GEL; Measurements: WH, MAP, NV, RH, LAK; Data analysis: WH, MAP; Supervision of work: SPJ, RDV; Manuscript writing: WH, MAP. All authors read and approved the manuscript. All authors had access to the data used in the study and accept responsibility for the decision to submit the manuscript for publication.

## References

1. Pizzolla A, Nguyen THO, Smith JM, Brooks AG, Kedzierska K, Heath WR, et al. Resident memory CD8 T cells in the upper respiratory tract prevent pulmonary influenza virus infection. Science Immunology. 2017;2(12).

2. Schenkel JM, Fraser KA, Beura LK, Pauken KE, Vezys V, Masopust D. Resident memory CD8 T cells trigger protective innate and adaptive immune responses. Science. 2014;346(6205):98–101.

3. Knisely JM, Buyon LE, Mandt R, Farkas R, Balasingam S, Bok K, et al. Mucosal vaccines for SARS-CoV-2: scientific gaps and opportunities-workshop report. NPJ Vaccines. 2023;8(1):53.

4. Lavelle EC, Ward RW. Mucosal vaccines - fortifying the frontiers. Nat Rev Immunol. 2022;22(4):236–50.

5. Lim JME, Ottolini S, Hang SK, Qui MD, Chia A, Low JGH, et al. Dynamics of virus-specific CD8+ T cells in the human nasal cavity. Mucosal Immunol. 2025.

6. Ramirez SI, Faraji F, Hills LB, Lopez PG, Goodwin B, Stacey HD, et al. Immunological memory diversity in the human upper airway. Nature. 2024;632(8025).

7. Lim JME, Tan AT, Bertoletti A. Protocol to detect antigen-specific nasal-resident T cells in humans. Star Protoc. 2023;4(1).

8. Roukens AHE, Pothast CR, König M, Huisman W, Dalebout T, Tak T, et al. Prolonged activation of nasal immune cell populations and development of tissue-resident SARS-CoV-2-specific CD8 T cell responses following COVID-19. Nature Immunology. 2022;23(1):23–+.

9. Jochems SP, Piddock K, Rylance J, Adler H, Carniel BF, Collins A, et al. Novel Analysis of Immune Cells from Nasal Microbiopsy Demonstrates Reliable, Reproducible Data for Immune Populations, and Superior Cytokine Detection Compared to Nasal Wash. Plos One. 2017;12(1).

10. Huisman W, Roex MCJ, Hageman L, Koster EAS, Veld SAJ, Hoogstraten C, et al. Tracking the progeny of adoptively transferred virus-specific T cells in patients posttransplant using TCR sequencing. Blood Adv. 2023;7(5):812–27.

11. da Silva Antunes R, Weiskopf D, Sidney J, Rubiro P, Peters B, Lindestam Arlehamn CS, et al. The MegaPool Approach to Characterize Adaptive CD4+ and CD8+ T Cell Responses. Curr Protoc. 2023;3(11):e934.

12. Lissina A, Ladell K, Skowera A, Clement M, Edwards E, Seggewiss R, et al. Protein kinase inhibitors substantially improve the physical detection of T-cells with peptide-MHC tetramers. J Immunol Methods. 2009;340(1):11–24.

13. Huisman W, Leboux DAT, van der Maarel LE, Hageman L, Amsen D, Falkenburg JHF, et al. Magnitude of Off-Target Allo-HLA Reactivity by Third-Party Donor-Derived Virus-Specific T Cells Is Dictated by HLA-Restriction. Front Immunol. 2021;12:630440.

14. Browne DJ, Miller CM, Doolan DL. Technical pitfalls when collecting, cryopreserving, thawing, and stimulating human T-cells. Front Immunol. 2024;15:1382192.

15. Clarkson BD, Johnson RK, Bingel C, Lothaller C, Howe CL. Preservation of antigen-specific responses in cryopreserved CD4(+) and CD8(+) T cells expanded with IL-2 and IL-7. J Transl Autoimmun. 2022;5:100173.

16. Massey CJ, Diaz Del Valle F, Abuzeid WM, Levy JM, Mueller S, Levine CG, et al. Sample collection for laboratory-based study of the nasal airway and sinuses: a research compendium. Int Forum Allergy Rhinol. 2020;10(3):303–13.

17. Urban BC, Goncalves ANA, Loukov D, Passos FM, Reine J, Gonzalez-Dias P, et al. Inflammation of the nasal mucosa is associated with susceptibility to experimental pneumococcal challenge in older adults. Mucosal Immunol. 2024;17(5):973–89.

18. Lindeboom RGH, Worlock KB, Dratva LM, Yoshida M, Scobie D, Wagstaffe HR, et al. Human SARS-CoV-2 challenge uncovers local and systemic response dynamics (vol 631, pg 189, 2024). Nature. 2024;632(8025):E3–E.

19. Lim JME, Tan AT, Le Bert N, Hang SK, Low JGH, Bertoletti A. SARS-CoV-2 breakthrough infection in vaccinees induces virus-specific nasal-resident CD8+ and CD4+ T cells of broad specificity. J Exp Med. 2022;219(10).

20. Arnous R, Arshad S, Sandgren K, Cunningham AL, Carnt N, White A. Tissue resident memory T cells inhabit the deep human conjunctiva. Sci Rep. 2022;12(1):6077.

21. Morton B, Burr S, Chikaonda T, Nsomba E, Manda-Taylor L, Henrion MYR, et al. A feasibility study of controlled human infection with Streptococcus pneumoniae in Malawi. EBioMedicine. 2021;72:103579.

22. Jochems SP, Marcon F, Carniel BF, Holloway M, Mitsi E, Smith E, et al. Inflammation induced by influenza virus impairs human innate immune control of pneumococcus. Nat Immunol. 2018;19(12):1299–308.

23. Linder A, Portmann K, Eyer K. The impact of cryopreservation on cytokine secretion and polyfunctionality in human PBMCs: a comparative study. Front Immunol. 2024;15:1478311.

24. Gallo O, Locatello LG, Mazzoni A, Novelli L, Annunziato F. The central role of the nasal microenvironment in the transmission, modulation, and clinical progression of SARS-CoV-2 infection. Mucosal Immunol. 2021;14(2):305–16.

25. Coates ML, Richoz N, Tuong ZK, Bowyer GS, Lee CYC, Ferdinand JR, et al. Temporal profiling of human lymphoid tissues reveals coordinated defense against viral challenge. Nat Immunol. 2025;26(2):215–29.

26. Dolton G, Lissina A, Skowera A, Ladell K, Tungatt K, Jones E, et al. Comparison of peptide-major histocompatibility complex tetramers and dextramers for the identification of antigen-specific T cells. Clin Exp Immunol. 2014;177(1):47–63.

27. Gao F, Mallajosyula V, Arunachalam PS, van der Ploeg K, Manohar M, Roltgen K, et al. Spheromers reveal robust T cell responses to the Pfizer/BioNTech vaccine and attenuated peripheral CD8(+) T cell responses post SARS-CoV-2 infection. Immunity. 2023;56(4):864–78 e4.

28. Bowyer G, Rampling T, Powlson J, Morter R, Wright D, Hill AVS, et al. Activation-induced Markers Detect Vaccine-Specific CD4(+) T Cell Responses Not Measured by Assays Conventionally Used in Clinical Trials. Vaccines (Basel). 2018;6(3).

29. Reiss S, Baxter AE, Cirelli KM, Dan JM, Morou A, Daigneault A, et al. Comparative analysis of activation induced marker (AIM) assays for sensitive identification of antigen-specific CD4 T cells. PLoS One. 2017;12(10):e0186998.

30. Maecker HT, Hassler J, Payne JK, Summers A, Comatas K, Ghanayem M, et al. Precision and linearity targets for validation of an IFNgamma ELISPOT, cytokine flow cytometry, and tetramer assay using CMV peptides. BMC Immunol. 2008;9:9.

31. Karlsson AC, Martin JN, Younger SR, Bredt BM, Epling L, Ronquillo R, et al. Comparison of the ELISPOT and cytokine flow cytometry assays for the enumeration of antigen-specific T cells. J Immunol Methods. 2003;283(1-2):141–53.

32. Binayke A, Zaheer A, Vishwakarma S, Singh S, Sharma P, Chandwaskar R, et al. A quest for universal anti-SARS-CoV-2 T cell assay: systematic review, meta-analysis, and experimental validation. NPJ Vaccines. 2024;9(1):3.

