## supplementary figures for "Systematic Evaluation of Nasal Immune Cell Sampling and Antigen-specific T cell Detection using Cryopreserved Nasal Swabs"

In order of manuscript text

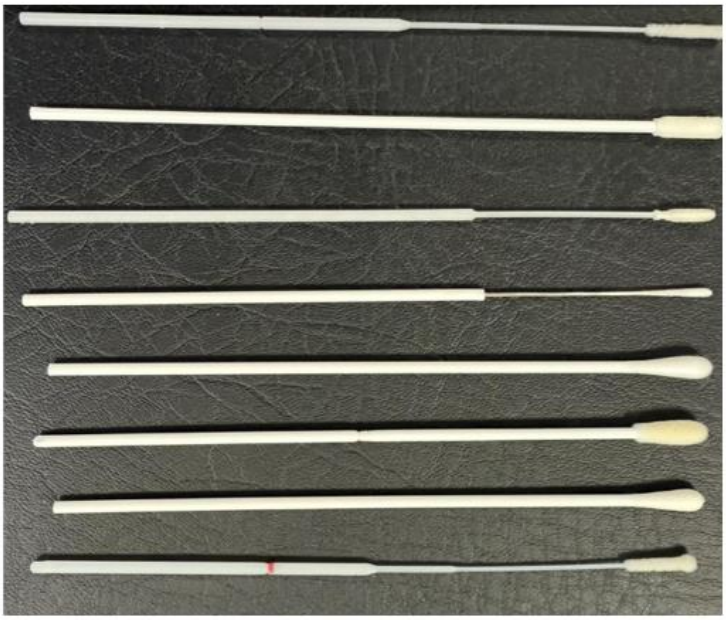

| | Flocked | Flex-tip |
|---|---|---|
| A | Yes | Yes |
| B | Yes | No |
| C | Yes | Yes |
| D | No | Yes |
| E | No | No |
| F | Yes | No |
| G | No | No |
| H | Yes | Yes |

**Supplementary Figure 1. Comparison of swab types used for inferior turbinate sampling.** Eight different swab types were evaluated for their cellular yield following swabbing of the inferior turbinate for 10 seconds. Swabs from two manufacturers (Puritan and Copan), including both flocked and flexible-tip designs, were compared. Cellular yield was assessed between different swabs.

A B

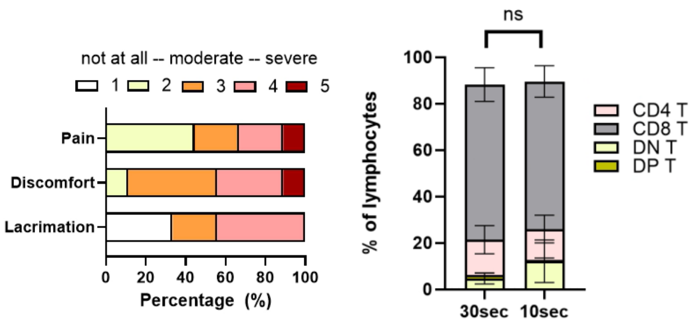


**Supplementary Figure 2. Tolerability and yield of flocked swabs per individual when nostrils are separately sampled**. **A**) General tolerability was assessed of 9 individuals where flocked swabs were collected from the inferior turbinate, randomized across left and right nostril. Flocked swabs were inserted for 10seconds. Lacrimation (Lac), discomfort (Dis) and pain were assessed. **B** Cellular composition of live-lymphocytes from freshly collected flocked swabs from the inferior turbinate were compared between 10 seconds and 30 seconds.

Statistical difference was assessed with Two-way Anova with Sidak's correction for multiple testing (B); * *P<0.05*, ****P < 0.001;

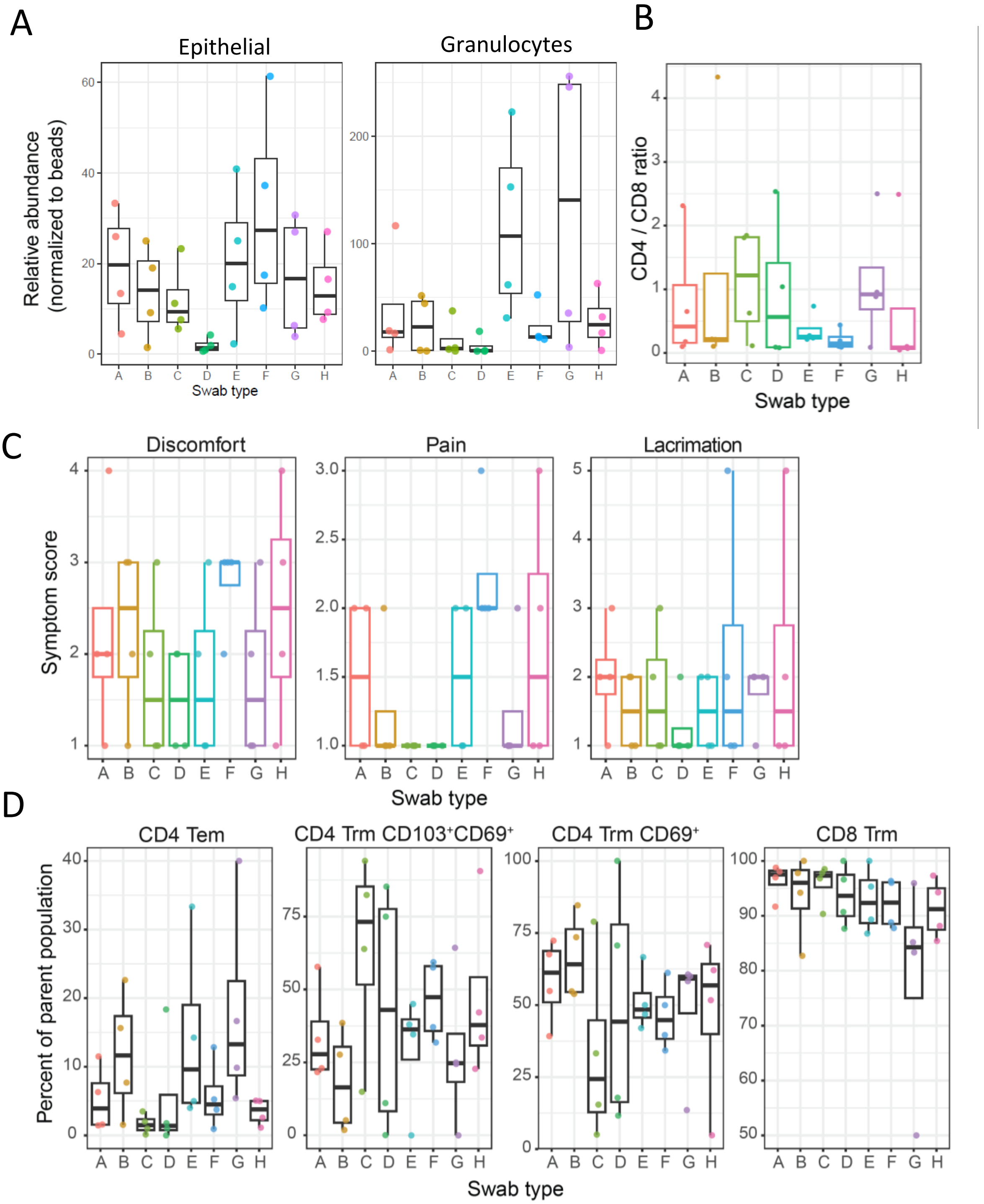


**Supplementary Figure 3. Tolerability and cellular yield of different swab types.** Eight swab types were evaluated for cellular yield and tolerability following 10 seconds of swabbing of the inferior turbinate. Swabs from two manufacturers (Puritan and Copan), including both flocked and non-flocked designs as well as swabs with flexible or rigid tips, were compared. Swab type H corresponded to the commonly used COPAN®503CS01. **A**) Yield of epithelial cells and granulocytes is shown for all eight swab types. **B**) CD4:CD8 T cell ratios are shown for each swab type. **C**) Tolerability was assessed by measuring perceived discomfort, pain, and lacrimation. **D**) Frequencies of tissue-resident memory (Trm) T cell subsets are shown for each swab type as percentage of parent population being either CD4 T cells or CD8 T cells.

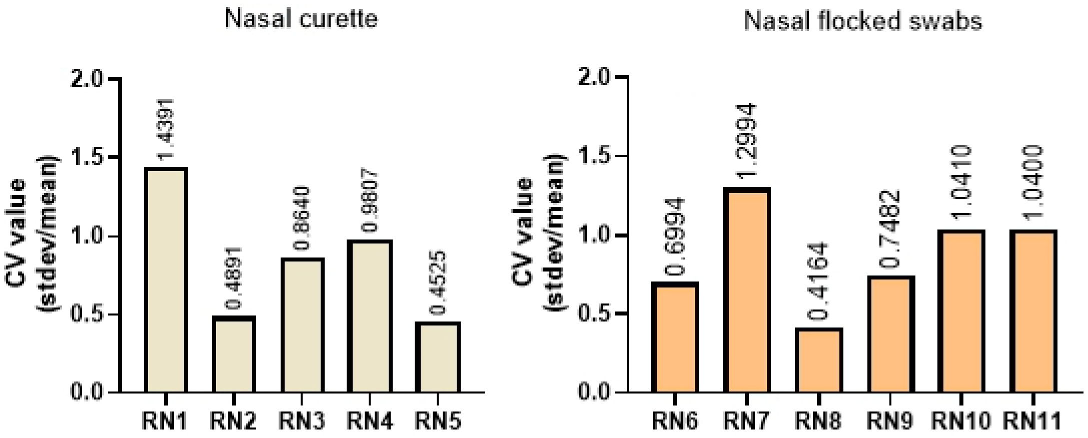


**Supplementary Figure 4. Variability between operators that collected nasal curretes or nasal flocked swabs**. Variability between operators in yield of lymphocytes was compared between using 2 nasal curretes (left;RhinoPro©) and one nasal flocked swabs (right; COPAN®). Coeficients of variations were calculated using standard deviation of the lymphocyte counts divided by the mean.

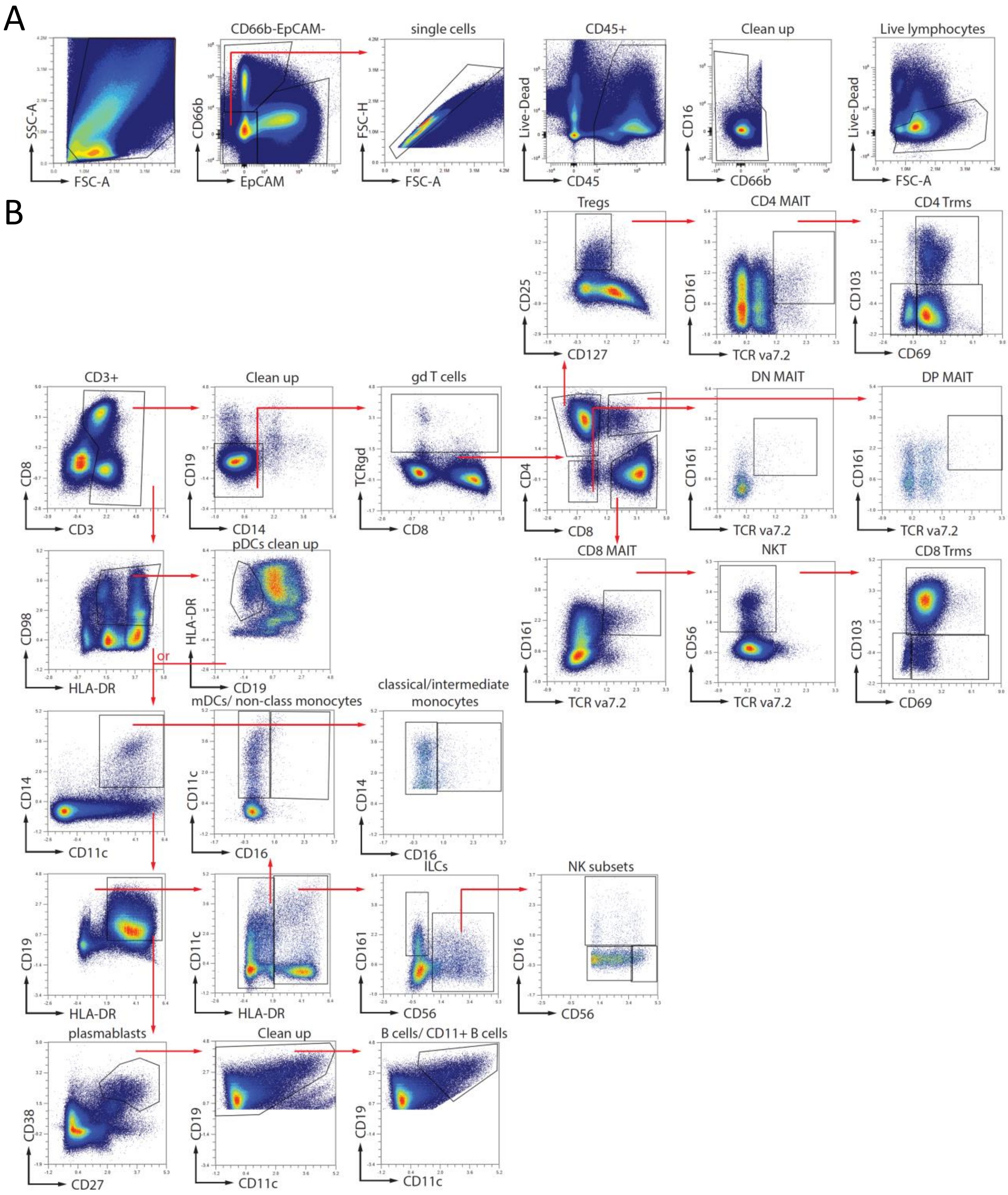


**Supplementary Figure 5. Gating strategy to define nasal-derived immune cell populations**. Concatenated flow cytometry plots are shown of 40 different samples taken from the Inferior, mid and superior turbinate and nasopharynx. Sampling was conducted by two independent operators. **A**) Gating strategy to retrieve granulocytes (CD66b+EpCAM-), epithelial cells (CD66b-EpCAM+) and lymphocytes (CD66b-EpCAM-). Live lymphocytes were further used to normalize all markers across the two experiments using FlowSOM k=3 with Cytonorm**. B**) live lymphocytes were further gated using normalized expression for all markers and started with gating CD3+ and CD3- immune cells.

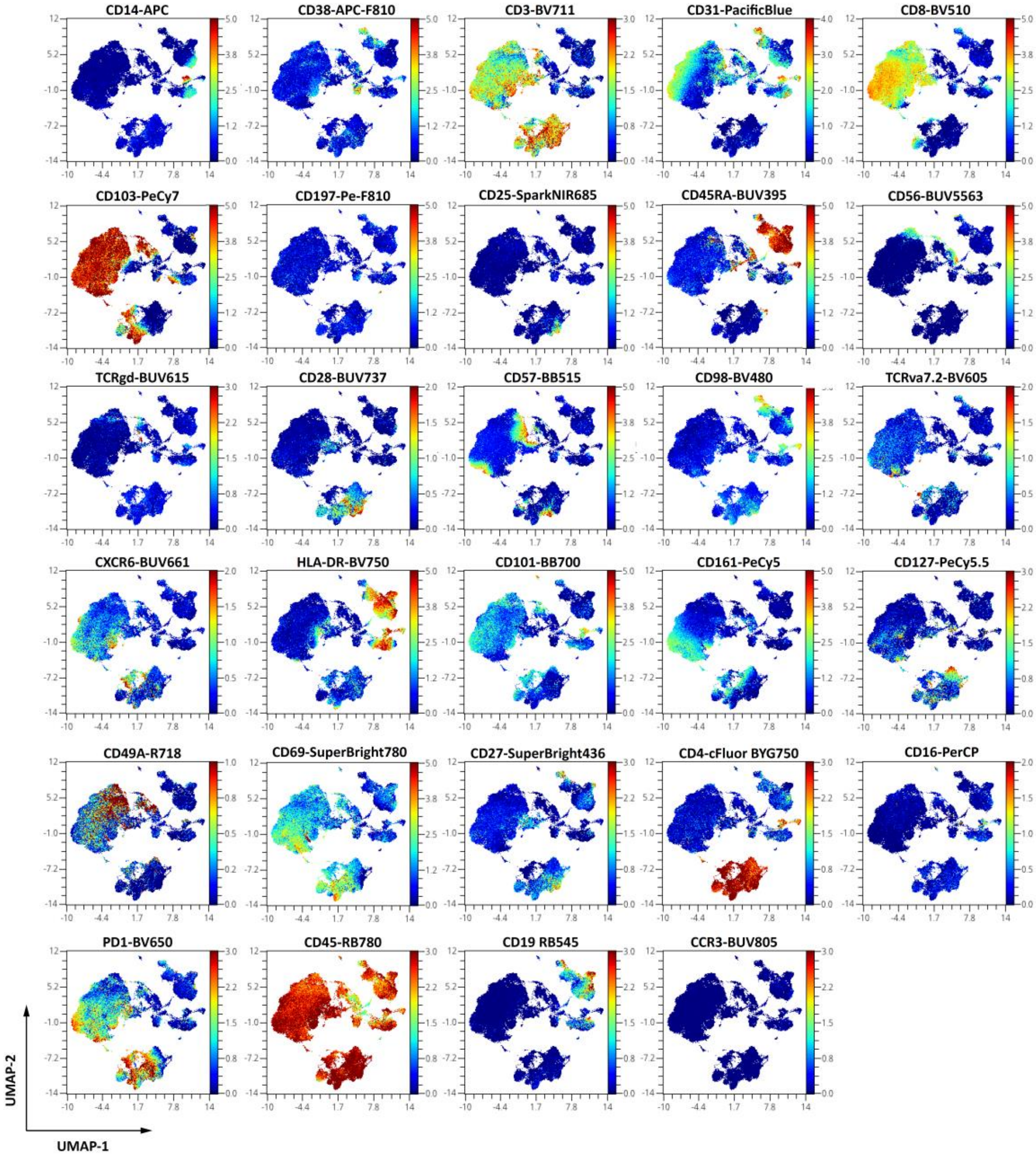


**Supplementary Figure 6. Overview of marker expression of nasal immune cells overlaid on UMAP.** UMAPs for 84k nasal cells, 21k cells per group, to cluster cells based on 29 different markers.

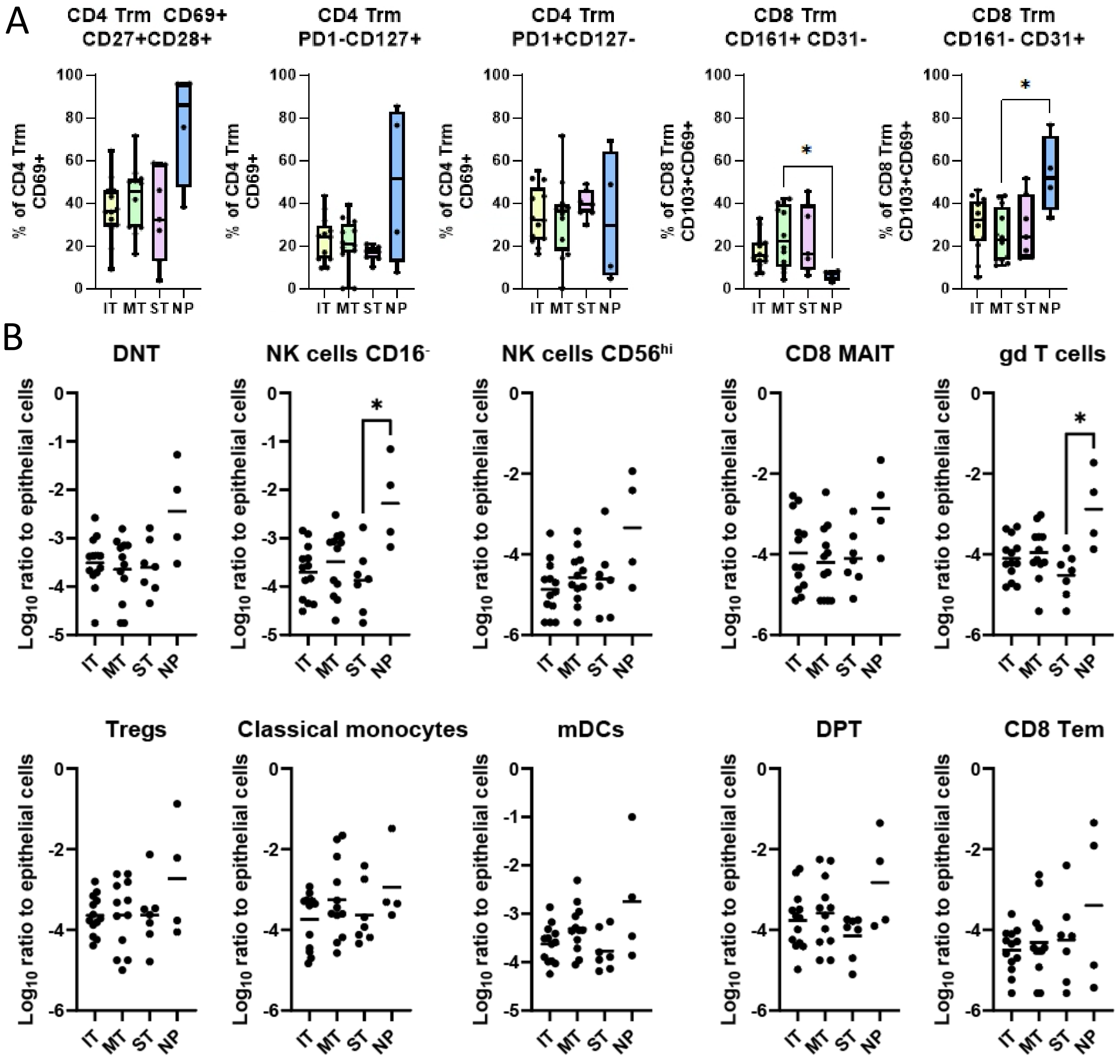


**Supplementary Figure 7**. **Distinct immune-cell phenotype and composition of nasal swabs from the upper respiratory tract.** Nasal swabbing using flocked swabs (COPAN) was performed at different anatomical locations (inferior turbinate; (IT) middle turbinate (MT), superior turbinate (ST) or nasopharynx (NP)) of the upper respiratory tract. Sampling was performed by two independent operators. **A)** Different subsets of tissue resident memory T cells are shown as percentage of CD4 Trms expressing CD69 or CD8 Trms expressing CD103 and CD69. **B)** Normalized abundances (log10 ratio relative to epithelial cells) of live nasal immune cell populations. Individual values and box plots are shown. Samples in which a given cell population was undetectable were assigned the lowest detectable value observed for that population.

Statistical differences were assessed using Kruskal Wallis tests with Dunn's correction for multiple testing (A and B). **P* < 0.05

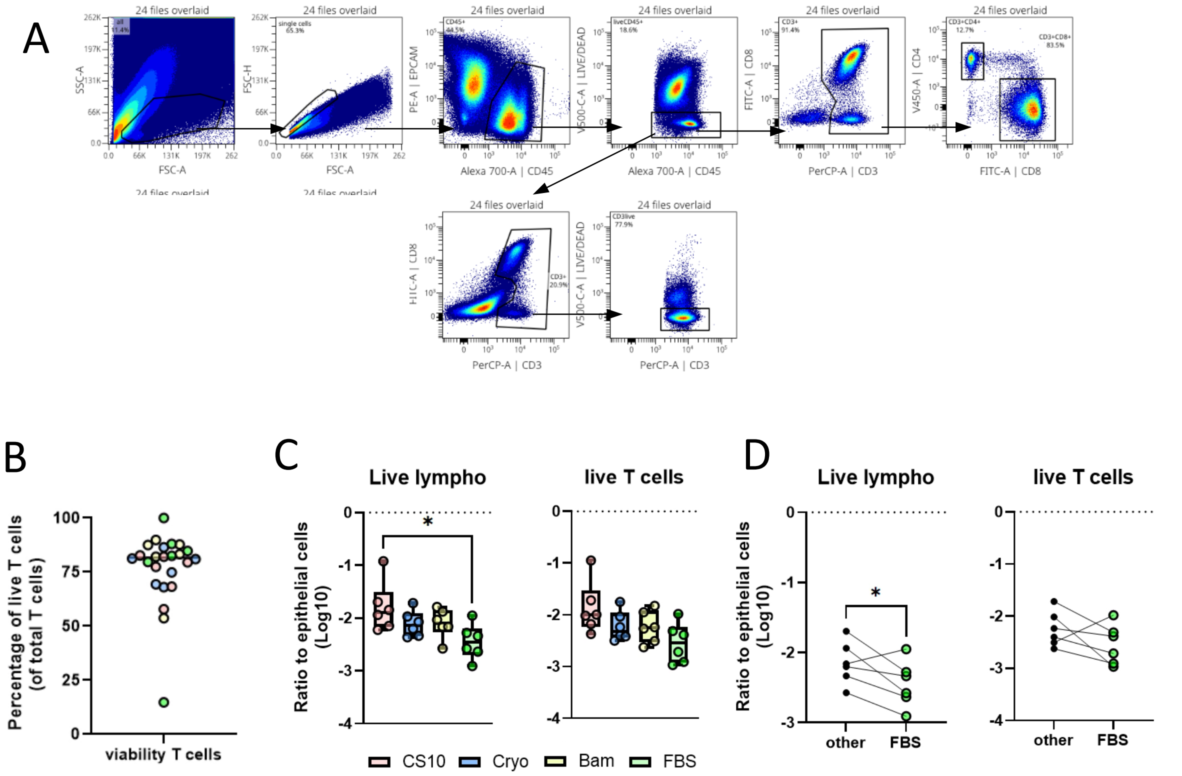


**Supplementary figure 8. Gating strategy and Comparison of different commercially available freezing media for the cryopreservation of nasal cells from flocked swabs.** Nasal swabbing using flocked swabs was performed at the inferior turbinate (IT) and immediately put into freeze media (Bambunker, CryoABC, Cryostor CS10 and 10%FBS/DMSO) and frozen according manufacturers protocol. In total, samples from 12 healthy donors, randomized across left and right nostril for different freeze media (n=6 each) were assessed. **A**) Shown is a gating strategy used to gate live lymphocytes and live T cells. **B**) Viability of T cells is shown in a dotplot. Samples are colored according to freezing media used. **C and D)** Ratio of live lymphocytes and live T cells normalized to the number of epithelial cells from the same sample. Individuals and box plots are shown where populations are shown as log10 transformed values (**B**) or paired samples are shown whereby Bambunker, CryoABC and Cryostor CS10 frozen samples are considered as other and compared to 10%FBS/DMSO (**C**).

Statistical differences were assessed with Kruskal Wallis test with Dunn's correction for multiple testing (B) or paired t tests (C). *$P < 0.05$;

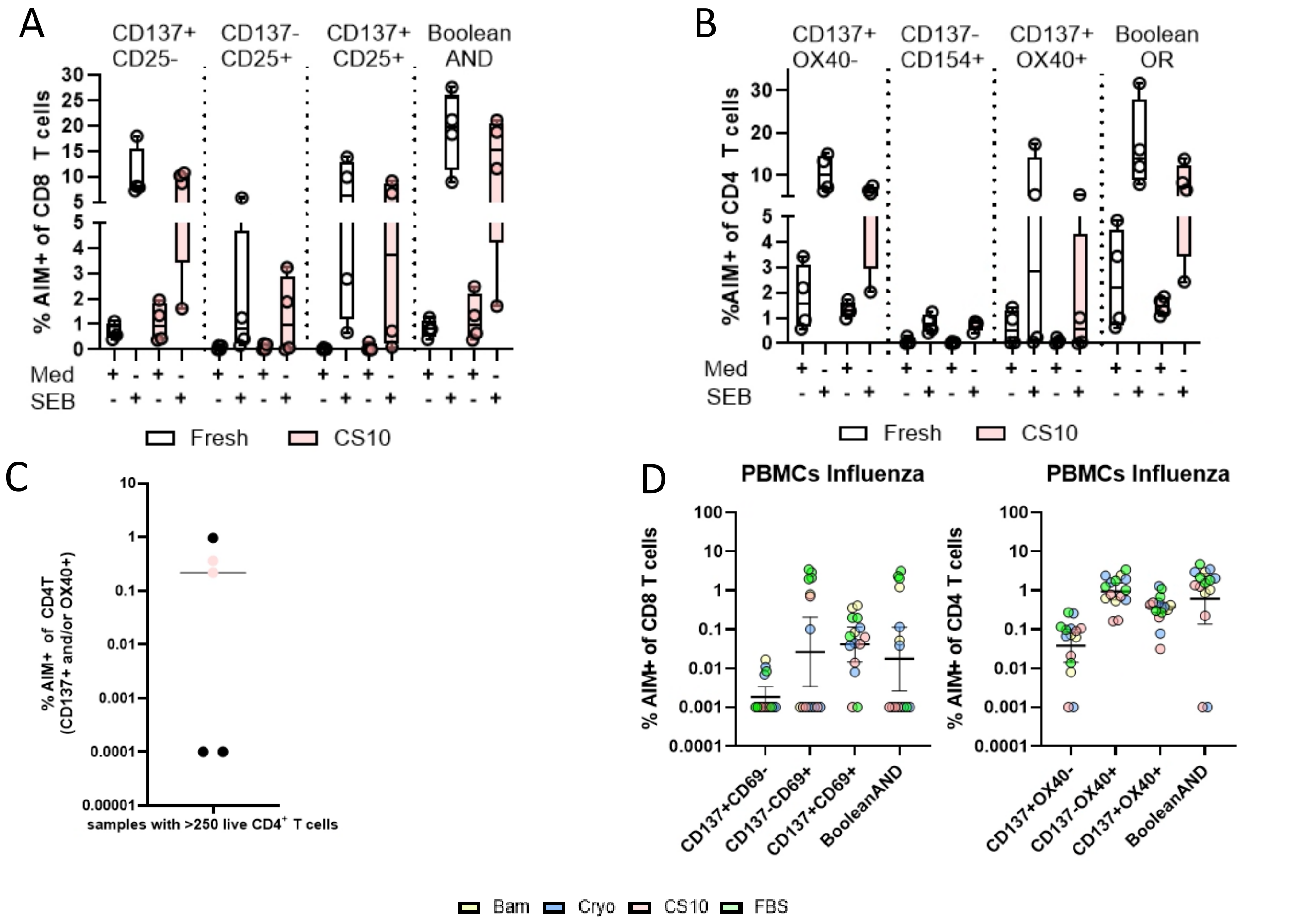


**Supplementary Figure 9. Activation induced marker (AIM) assay to detect Influenza-specific and SEB-reactive T cells using cryopreserved nasal flocked swabs and PBMCs.** Nasal swabbing using FLOQswabs was performed at the inferior turbinate (IT) and samples (n=4) from both nostrils were mixed and 50% was first frozen and then thawed, processed and stimulated with SEB while the other 50% was immediately stimulated with SEB. Additionally, paired nasal flocked samples and PBMCs cryopreserved using different freeze media (n=4 each) from 12 healthy donors were similarly processed and used to study Influenza-specific T cells. **A and B**) Activated CD8 (A) or CD4 (B) T cells that were stimulated with SEB immediately after collection (white) or after cryopreservation with cryostor CS10 (pink). Percentage of cells that show activation induced markers (AIM) was assessed and are shown for each combination separately. AIM+ represents a Boolean AND gate for all combinations shown. **C**) Shown are dotplots of activated CD4$^{+}$ T cells that were stimulated fresh (black) or after cryopreservation using CS10 (pink) with an overlapping pool of Influenza peptides (non-HA). Samples with more than 250 live CD4$^{+}$ T cells were only analysed. Data shown are medium (DMSO) subtracted. Percentage of AIM+ CD4$^{+}$ T cells represent the sum of cells that express either CD137 and/or OX40. Samples without AIM+ events were plotted at 0.0001. **D**) Activated CD8 or CD4 T cells from PBMCs that were stimulated with immunodominant peptides from non-HA proteins from Influenza after cryopreservation with different freezing media.

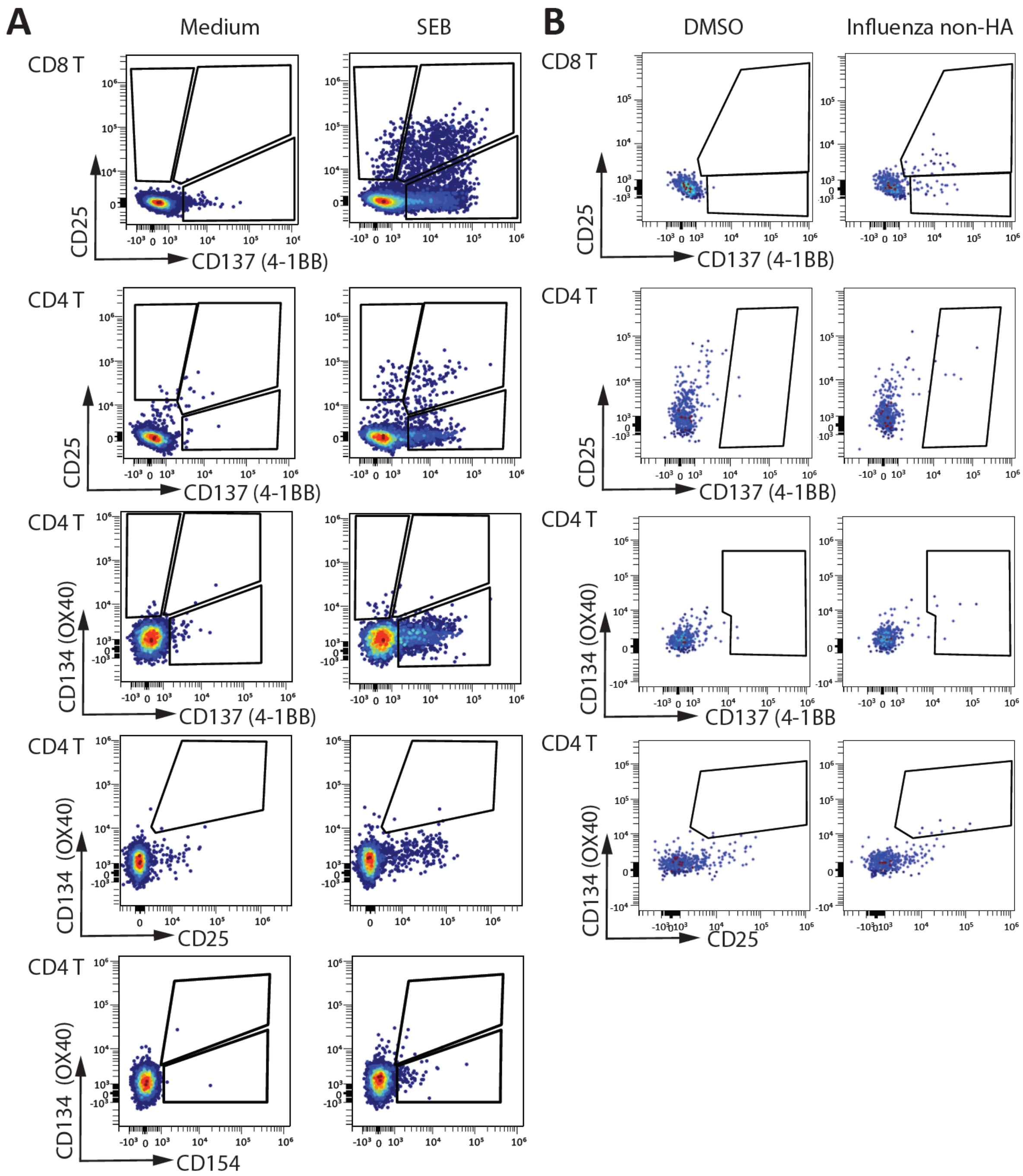


**Supplementary Figure 10. Gating strategy for activation induced marker (AIM) assay**. **A and B**) Nasal derived immune cells were stimulated overnight for 16 hours with (**A**) medium or SEB and (**B**) DMSO or overlapping peptide pools covering non-HA proteins of influenza. To quantify percentages of activation induced markers (AIM) of CD4 and CD8 T cells after stimulation, nasal immune cells were first gated on Live-lymphocytes excluding granulocytes and epithelial cells. Subsequently, CD3+ T cells were gated and CD4 and CD8 T cells were assessed separately for AIM expression. Percentages of cells that show AIM+ were assessed and are shown for each combination separately.

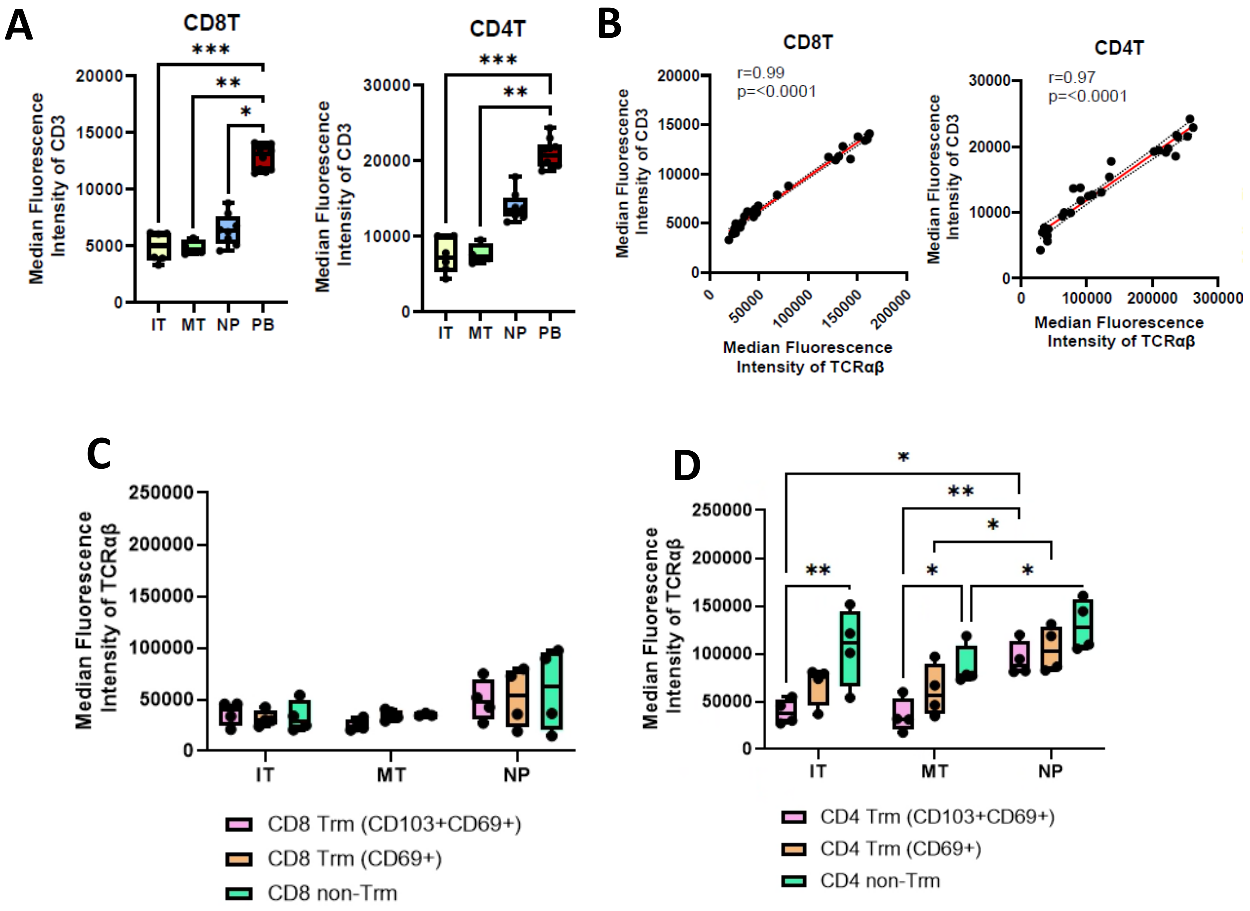


**Supplementary Figure 11. Intrinsic differences of the TCR:CD3 complex of nasal and peripheral T cells**. **A**) A) Median Fluorescent Intensity of CD3 are shown of CD4 and CD8+ T cells from PBMCs (red) and different anatomical locations of the upper-respiratory tract, obtained using flocked swabs. B) Pearson correlation plots for the median fluorescence intensity of CD3 against the median fluorescence intensity of TCRαβ are shown for CD8 and CD4 T cells. C and D) Median Fluorescent Intensity of TCRαβ are shown of different subsets of tissue resident memory T cells and T cells that lack tissue-resident markers and compared across samples taken at different anatomical locations of the upper respiratory tract.

Statistical differences were assessed with Kruskal Wallis test with Dunn's correction for multiple testing (A, C and D). *$P < 0.05$; **$P < 0.01$; ***$P < 0.001$
